# Genome-Wide Association Study of Over 427,000 Individuals Establishes Executive Functioning as a Neurocognitive Basis of Psychiatric Disorders Influenced by GABAergic Processes

**DOI:** 10.1101/674515

**Authors:** Alexander S. Hatoum, Claire L. Morrison, Evann C. Mitchell, Max Lam, Chelsie E. Benca-Bachman, Andrew E. Reineberg, Rohan H. C. Palmer, Luke M. Evans, Matthew C. Keller, Naomi P. Friedman

**Affiliations:** Institute for Behavioral Genetics, University of Colorado Boulder; Department of Psychology and Neuroscience, University of Colorado Boulder; Department of Psychiatry, University of Washington St. Louis Medical School; University of Texas Southwestern Medical School; Division of Psychiatry Research, The Zucker Hillside Hospital, Glen Oaks, NY, 11004, USA; Stanley Center for Psychiatric Research, Broad Institute of Harvard and MIT, Cambridge, MA, 02142, USA; The Behavioral Genetics of Addiction Laboratory, Department of Psychology, Emory University, Atlanta, GA, USA; Department of Ecology and Evolutionary Biology, University of Colorado Boulder

**Keywords:** Executive Functioning, Neurocognitive Functioning, Genome-wide Association Analysis, Genetic Correlations, Latent variable measurement

## Abstract

Deficits in executive functions (EFs), cognitive processes that control goal-directed behaviors, are associated with psychopathology and neurological disorders. Little is known about the molecular bases of EF individual differences; existing EF genome-wide association studies (GWAS) used small sample sizes and/or focused on individual tasks that are imprecise measures of EF. We conducted a GWAS of a Common EF (cEF) factor based on multiple tasks in the UK Biobank (*N*=427,037 European-descent individuals), finding 129 independent genome-wide significant lead variants in 112 distinct loci. cEF was associated with fast synaptic transmission processes (synaptic, potassium channel, and GABA pathways) in gene-based analyses. cEF was genetically correlated with measures of intelligence (IQ) and cognitive processing speed, but cEF and IQ showed differential genetic associations with psychiatric disorders and educational attainment. Results suggest that cEF is a genetically distinct cognitive construct that is particularly relevant to understanding the genetic variance in psychiatric disorders.

## Introduction

Deficits in executive functions (EFs), higher-level cognitive abilities that enable control over thoughts and actions during goal-directed behavior^1^, are debilitating for daily life and are a hallmark of brain disorders. In particular, EF deficits are associated with almost all psychiatric disorders, leading some to suggest that EF deficits are a risk factor for general psychopathology (i.e., the *p* factor)^2–5^. Recent work using single nucleotide polymorphism (SNP) effects from large genome-wide associations studies (GWAS) to estimate genetic correlations suggests that cognition–psychopathology associations may be partially genetic in origin^6,7^. These studies have focused on general cognitive ability or intelligence quotient (IQ), the cognitive construct with the largest GWAS sample sizes. Within the GWAS literature, there is an implicit assumption that IQ captures most of the genetic variance across cognitive phenotypes. However, adult phenotypic and twin studies suggest that a Common EF (cEF) factor capturing variance shared across diverse EF tasks is distinguishable from general IQ at the phenotypic and genetic levels, and predicts behavior over and above IQ^1,8,9^. Here, we conduct a large GWAS of a cEF factor score generated from data in the UK Biobank (UKB) study^10^ to discover cEF’s molecular underpinnings. We then use the GWAS results to test the hypotheses that cEF is genetically separable from IQ and cognitive processing speed, and that it is the cognitive dimension most relevant for understanding genetic variation underlying psychopathology.

EF is a blanket term for a family of cognitive functions^11^. Commonly studied EFs include response inhibition, interference control, working memory updating and capacity, and mental set shifting^1^. EFs are typically measured with laboratory cognitive tasks such as the antisaccade, Stroop, *n*-back, complex working memory span, and task-switching paradigms, or with neuropsychological tests such as trail making form B, digit span, and Wisconsin card sorting test. Because EFs are control processes, an EF task requires the combination of lower-level cognitive processes (e.g., visual processing of stimuli) in addition to the higher-level EF processes of interest (e.g., biasing of attention towards task-relevant lower-level information)^12^. These lower-level processes contribute to individual differences in performance on specific tasks, leading to the “task impurity problem”^12^. This task impurity means that GWAS loci and molecular processes associated with individual EF tasks may capture cognitive processes other than cEF. Individual EF tasks can also show low reliability^12^, decreasing power for association tests.

The task impurity problem is alleviated by extracting common variance across multiple EF tasks with a cEF factor^8,13,14^. Four independent twin studies showed that across samples and ages, cEF is moderately to highly heritable^13–15^ (46% to 100%) and highly phenotypically and genetically stable across time^9,16^. However, little is known about the molecular underpinnings of cEF. Most historical perspectives from the candidate gene^17^ and animal^18^ literature have argued that neurocognitive function is supported by metabotropic processes, in particular the slow neuromodulator effects of the dopaminergic systems. However, recent work in humans with ketamine suggests that fast ionotropic processes influence neurocognitive ability, in particular, the excitatory neurotransmitter glutamate (via activation of Anti-N-methyl-D-aspartate (NMDA) receptors)^19^. Fast inhibitory GABAergic processes have also been studied in relation to EFs, but are often neglected in the literature^20^.

Existing GWAS of EF have had insufficient power to test hypotheses regarding the molecular mechanisms that underlie EF. To date, the largest GWAS of neurocognitive tasks included 1,311 to 32,070 individuals, depending on the task, and found a single genome-wide significant association for a processing speed task^21^. If cEF follows the pattern observed for almost all other complex traits, it is likely that larger samples will be required to discover and differentiate the molecular pathways associated with cEF. Furthermore, all previous molecular genetic studies have measured EF using individual tasks. Using a factor score should both bolster power by increasing the effect sizes of SNP associations with cEF^12^ and reduce GWAS associations reflecting task impurity.

Because cEF reflects variance shared among multiple cognitive tasks, a natural question is whether cEF is synonymous with general cognitive ability or IQ. Data from several independent twin studies suggest that phenotypically and genetically, cEF is moderately to highly correlated with IQ (*r*=.53–.91; *r*G=.57–1.0)^8,13,22^. In adult samples^8,13^, cEF correlations with IQ are moderate (*r*=.53–.68; *r*G=.57–.59) and significantly lower than 1. Moreover, IQ genetically correlates with variance specific to working memory processes in addition to cEF^8,13^, suggesting that IQ variation is supported by both cEF and working memory-specific abilities in adults. Phenotypic literature also suggests that EFs show discriminant predictive validity of behavioral problems when controlling for IQ^23^. Genetic correlations derived from GWAS provide a new opportunity to evaluate whether cEF may capture distinct genetic variance from IQ and show stronger relations to psychopathology, including a *p* factor^3^ that would be impossible to evaluate in phenotypic/twin studies.

This study is the first GWAS of cEF using a factor based on multiple cognitive tasks. We generated a cEF factor score in the UKB sample of over 427,000 individuals of European ancestry based on the commonality of five EF tasks across multiple measurement occasions. We also estimated factor scores for IQ (verbal-numerical reasoning; *n*=216,381 in the genetic analysis) and cognitive processing speed (*n*=432,297 in the genetic analysis) for comparison. To further validate these factors, we computed polygenic scores (PGSs) for cEF and IQ based on these GWAS and tested whether they differentially predicted multiple EF latent variables and IQ in samples that were deeply phenotyped for these constructs in young adulthood.

We hypothesized that there would be a sizeable genetic correlation of cEF with IQ, but that this genetic correlation would be significantly less than 1.0. Moreover, we expected that cEF would show stronger genetic correlations than IQ or speed with psychopathology, and would predict a *p* factor when controlling for IQ and speed.

## Results

### GWAS of cEF Factor

Using confirmatory factor analysis, we calculated a cEF factor score in the UKB sample of 427,037 individuals (the “full” sample) and conducted a GWAS on this score as our main analysis (see Figure 1). The actual *n* for each EF task varied because individuals completed a different number of online tasks (see Table 1 for task descriptive statistics and Table 2 for *r*Gs among tasks). We tested consistency by conducting GWAS in two UKB subsamples that were densely (*n*=93,024) or only sparsely phenotyped (*n*=256,135 after removing relatives of those in the densely phenotyped sample).

**Figure 1.**
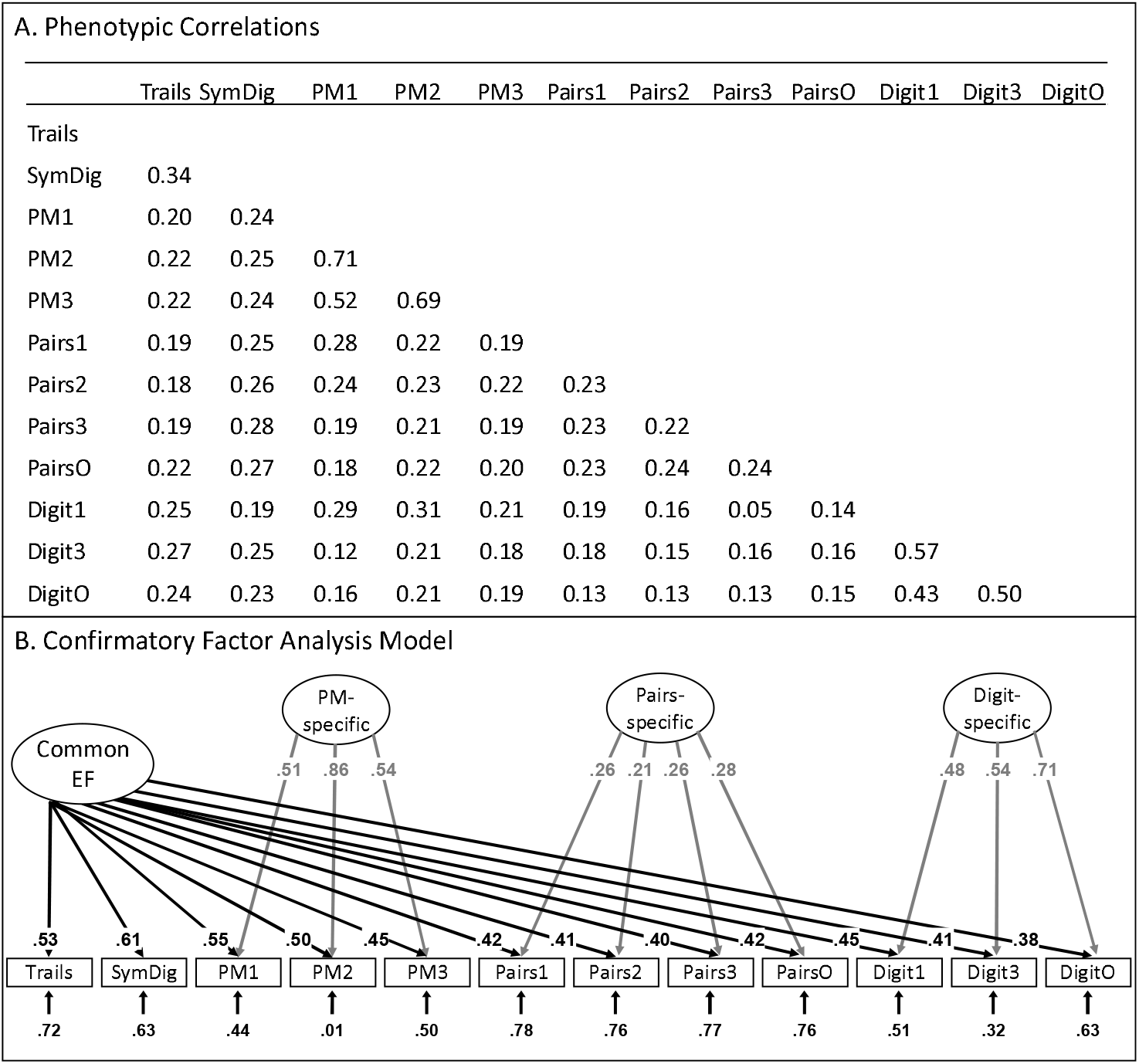
Development of a Common Executive Functioning (cEF) factor across cognitive tasks in the UK Biobank: (A) Correlations taken from Mplus; (B) Confirmatory factor analysis model used to extract factor scores. Ellipses indicate latent variables; rectangles indicate observed variables. Numbers on arrows are standardized factor loadings, and numbers at the end of arrows are residual variances. All parameters were statistically significant (*p* < .05). Trails= trail making (online); SymDig= symbol-digit substitution (online); PM= prospective memory; Pairs= pairs memory; Digit= digit span. Task names with 1= first assessment; with 2= repeat assessment; with 3= imaging visit assessment; with O= online follow-up. Directionality was reversed for some variables so that for all variables, higher scores indicate better performance.

**Table 1.** Descriptive statistics for cognitive measures used to obtain factor scores

| Measure | *N* | Mean | SD | Min | Max | Skewness | Kurtosis |
| --- | --- | --- | --- | --- | --- | --- | --- |
| **Trail making** |  |  |  |  |  |  |  |
| Online<sup>a</sup> | 104,050 | 0.00 | 0.11 | -0.44 | 0.44 | 0.48 | 0.73 |
| Numeric<sup>b</sup> | 104,052 | 1.57 | 0.14 | 1.14 | 2.87 | 0.65 | 0.67 |
| Alphanumeric<sup>b</sup> | 104,050 | 1.80 | 0.15 | 1.31 | 2.87 | 0.49 | 0.46 |
| **Symbol-digit substitution** |  |  |  |  |  |  |  |
| Online | 117,785 | 19.76 | 5.11 | 0 | 40 | -0.40 | 0.54 |
| **Prospective Memory<sup>c</sup>** |  |  |  |  |  |  |  |
| Initial visit | 171,309 | 0.24 | -- | -- | -- | -- | -- |
| Repeat visit | 20,314 | 0.15 | -- | -- | -- | -- | -- |
| Imaging visit | 15,880 | 0.12 | -- | -- | -- | -- | -- |
| **Pairs Matching<sup>d</sup>** |  |  |  |  |  |  |  |
| Initial visit | 484,340 | 0.76 | 0.37 | 0.00 | 2.22 | 0.39 | 0.56 |
| Repeat visit | 20,085 | 0.70 | 0.34 | 0.00 | 2.06 | 0.33 | 0.55 |
| Imaging visit | 15,472 | 0.66 | 0.33 | 0.00 | 2.00 | 0.35 | 0.61 |
| Online | 114,828 | 0.83 | 0.37 | 0.00 | 2.31 | 0.39 | 0.26 |
| **Digit Span** |  |  |  |  |  |  |  |
| Initial visit | 50,116 | 6.69 | 1.34 | 2 | 12 | -0.32 | 0.84 |
| Imaging visit | 4,237 | 6.80 | 1.24 | 2 | 11 | -0.20 | 0.68 |
| Online | 111,086 | 6.92 | 1.49 | 2 | 11 | -0.38 | 1.09 |

**Note.** Descriptive statistics and sample information for each task loading on the common executive functioning (cEF) factor from the UKBiobank sample.
<sup>a</sup>Unstandardized residual of the log10-transformed alphanumeric path time after regressing out the log10-transformed numeric path time; only this score was used in the model.
<sup>b</sup>Log10-transformed total times in seconds to complete the numeric and alphanumeric paths; these variables were not used in the confirmatory factor analysis model but were used to obtain the residualized trails measure used in the model.
<sup>c</sup>Categorical variable coded as 1 for correct and 0 for incorrect on first try. The mean described proportion correct. Dashes indicate that other descriptive statistics were not calculated.
<sup>d</sup>Sum of the log10-transformed number of incorrect matches +1 in the 6- and 12-card rounds.

**Table 2.** Heritability (diagonal) and genetic correlations (off-diagonal) between Common Executive Functioning (cEF) indicators and cEF factor scores

| Measure | Symbol Digit | Pairs Memory | Digit Span | Prospect. Memory | Trail Making | Dense cEF | Sparse cEF | Full cEF |
| --- | --- | --- | --- | --- | --- | --- | --- | --- |
| **Symbol–Digit** | **0.1245**<br>**(0.0079)** |  |  |  |  |  |  |  |
| **Pairs Memory** | 0.6603<br>(0.0271) | **0.0713**<br>**(0.003)** |  |  |  |  |  |  |
| **Digit Span** | 0.3226<br>(0.0345) | 0.4420<br>(0.0263) | **0.1337**<br>**(0.0069)** |  |  |  |  |  |
| **Prospective Memory** | 0.4479<br>(0.0414) | 0.5982<br>(0.0348) | 0.4539<br>(0.0355) | **0.0527**<br>**(0.0039)** |  |  |  |  |
| **Trail Making** | 0.7126<br>(0.0322) | 0.7085<br>(0.0317) | 0.6530<br>(0.0293) | 0.5927<br>(0.0463) | **0.1136**<br>**(0.0084)** |  |  |  |
| **Dense sample cEF** | 0.8428<br>(0.0138) | 0.8580<br>(0.0207) | 0.6653<br>(0.0214) | 0.6416<br>(0.0365) | 0.9274<br>(0.0133) | **0.1894**<br>**(0.0105)** |  |  |
| **Sparse sample cEF** | 0.7031<br>(0.0307) | 0.9831<br>(0.0074) | 0.5580<br>(0.0259) | 0.7052<br>(0.0308) | 0.7771<br>(0.0381) | 0.9230<br>(0.0286) | **0.0696**<br>**(0.0038)** |  |
| **Full sample cEF** | 0.7683<br>(0.0178) | 0.9527<br>(0.0047) | 0.6164<br>(0.0178) | 0.7046<br>(0.0255) | 0.8452<br>(0.0215) | 0.9629<br>(0.0106) | 0.9892<br>(0.0073) | **0.0906**<br>**(0.0038)** |

**Note.** The heritability of each measure (standard error) is shown on the diagonal in bold-face type. The lower diagonal contains the genetic correlations of each indicator and common executive functioning (cEF) factor scores in the densely phenotyped (dense), sparsely phenotyped (sparse), and full samples, as estimated by LD score regression. When there were multiple assessments of the same task (pairs memory, digit span), the measure is the average of the $z$ -scores for all assessments, except for the categorical prospective memory task, for which the measure used for this table is the first assessment.

SNP-heritability of cEF estimated via BOLT-REML^24^ was 0.104 (se=0.002). We found 129 lead (r^2^ <.1) and 299 independent SNPs (r^2^ < .6) in 112 distinct loci that were significantly associated with cEF in the full sample, using BOLT^24^ to run a linear mixed model test of association controlling for age, age, sex, the first 20 principal components (PCs), and batch and site (Figure 2, supplemental Figures S1-S3, and supplemental Tables S1-S6). The most significantly associated SNP (rs12707117, β= −0.012, *p*=2.1e-26) is an expression Quantitative Trait Loci (eQTL) in cerebellar tissue mapped to *EXOC4*. Q-Q plots (supplemental Figure S1) showed departure from expected *p*-values under the null hypothesis for the full sample and subsamples (λ_full_=1.6946, λ_dense_=1.311, λ_sparse_=1.3101), but the low linkage disequilibrium (LD) score regression intercepts (full=1.0381, dense=1.0128, sparse=1.0238) suggest that this inflation reflects polygenicity rather than confounding stratification.

**Figure 2.**
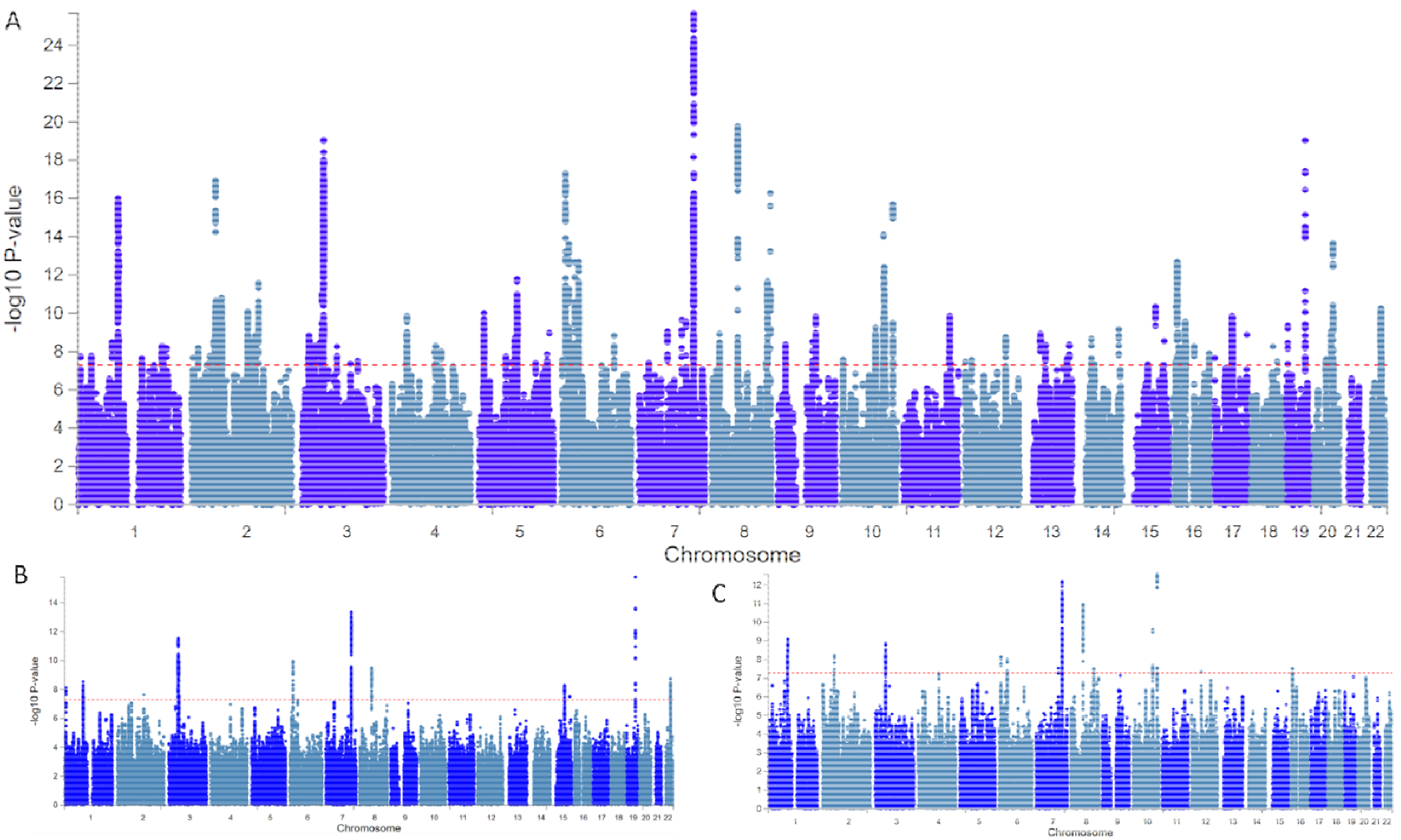
Manhattan plots for GWAS of Common Executive Functioning (cEF) in the full sample (Panel A), the densely phenotyped sample (Panel B), and the sparsely phenotyped sample (Panel C). Each dot is a single nucleotide polymorphism (SNP), chromosomes are organized on the x-axis, and the y-axis represents the negative log10 of the *p*-value for each SNP.

As shown in Table 2, the SNP-heritability for the densely-phenotyped sample (SNP-*h*^2^=0.189, se=0.011) was higher than for the sparsely-phenotyped sample (SNP-*h*^2^=0.070, se=0.004), as would be expected given that factor scores based on more tasks (densely-phenotyped) should have lower error variance (see online methods). However, the genetic correlation of the densely and sparsely phenotyped subsamples confirmed they measured substantially overlapping constructs (*r*G=0.923, se=0.029).

As expected given the smaller sample sizes, the subsamples showed weaker genome wide-discovery compared to the full sample. Yet despite the three-fold smaller sample size in the densely phenotyped sample, we identified a similar number of genome-wide significant loci in both samples: 34 independent and 15 lead SNPs in the densely phenotyped sample; 30 independent and 16 lead SNPs in the sparsely phenotyped sample (see supplemental Table S7). The fact that we detected as many SNPs in the smaller densely phenotyped sample as we did in the larger sparsely phenotyped sample suggests we had greater measurement precision in the densely phenotyped subsample, and that adequate measurement of phenotypes is an important aspect of discovering cEF-associated loci. However, the larger number of identified loci when using the combined dataset demonstrates the statistical power gained from utilizing our factor-based approach to leverage the entire sample. Therefore, the following analyses use the full sample.

### Genetic Separability of cEF and IQ

#### Genetic Correlation

To assess the genetic separability of cEF and IQ, we estimated their genetic correlation using LD Score regression (LDSC) and BOLT-REML. We first conducted a GWAS of IQ and Speed factor scores using relevant measures from the UKB. We did so because previously published GWAS of general cognitive ability and IQ included measures of EF tasks^6,7^, which might confound the test whether they are separable constructs. cEF factor scores were phenotypically correlated with IQ factor scores (*r*=.35, *p*<.001) and Speed factor scores (*r*=.28, *p*<.001); IQ and Speed factor scores were weakly correlated with each other (*r*=.17, *p*<.001), demonstrating divergence at the phenotypic level. Based on the GWAS association statistics using LDSC, we estimated the genetic correlation between cEF and IQ at .743 (se=.013, *p*=1.00e-221), which was significantly lower than 1 (*p*=1.4e-59). Similarly, using BOLT-REML we estimated the genetic correlation to be .766 (se=.007), *p*<1e-300); the 95% confidence interval (.752–.778) did not include 1.

These SNP-based genetic correlations reflect the genetic separability of cEF and IQ and are similar to those from twin-based *r*G estimates of IQ and cEF (*r*G=.69) in middle aged adults^13^. The genetic correlations are higher than the phenotypic correlations because genetic correlations consider the covariance only of the reliable genetic variance (standardized by the genetic variances of both traits), whereas phenotypic correlation consider the total covariance adjusted for the total phenotypic variances, which can include unique environmental effects and measurement error. The latter thus have a proportionally larger denominator, leading to smaller phenotypic than genetic correlations. Genetic correlations of our IQ factor score and past GWAS IQ^6,7^ were close to unity (*r*Gs = .964–.982; supplemental Table S8).

#### GWAS of cEF conditioned on IQ

Due to cEF’s high genetic correlation with IQ in particular, we used mt-COJO^25^ to estimate unique genetic effects associated with cEF and IQ, when conditioned on each other (supplemental Figure S4, supplemental Table S9). Due to the moderate to high genetic correlation between the two constructs, we anticipated that statistical power would be lower for this conditional GWAS. Consistent with this expectation, we identified 41 lead SNPs significantly associated with cEF when conditioned on IQ. Notably, the EXOC4 variant remained significantly associated with cEF, as did APOE. We identified 17 lead SNPs significantly associated with IQ, conditioning on cEF (supplemental Table S10). These results indicate that there are cEF-specific genetic effects when controlling for IQ, and IQ-specific genetic effects, when controlling for cEF, further demonstrating their separability.

#### Polygenic score analyses

To further validate the cEF phenotype in UKB and confirm that our cEF SNPs showed some commonality across ages, we created polygenic scores (PGS) of cEF and IQ in two young adult twin samples that were deeply phenotyped on multiple EF latent variables (cEF, Updating-specific, and Shifting-specific) and full-scale IQ. To maximize power and minimize the number of tests, we created the model shown in Figure 3. This Unity/Diversity EF model^1^ integrates full-scale IQ data and the multiple waves of EF data in line with our previously published twin models of these data^15,16^. All measures were residualized on age, sex, and age*sex within each sample at each wave prior to analysis, and the model included 10 PCs and batch as covariates, as shown. We restricted the PGS analysis to individuals of European ancestry (based the first 3 PCs), resulting in a final *N* of 916 (results for less conservative ancestry restrictions are also shown in supplemental Table S11). The variance explained by these PRS are expected to be substantially less than the SNP-*h*^2^ because the discovery sample size is finite^26^.

**Figure 3.**
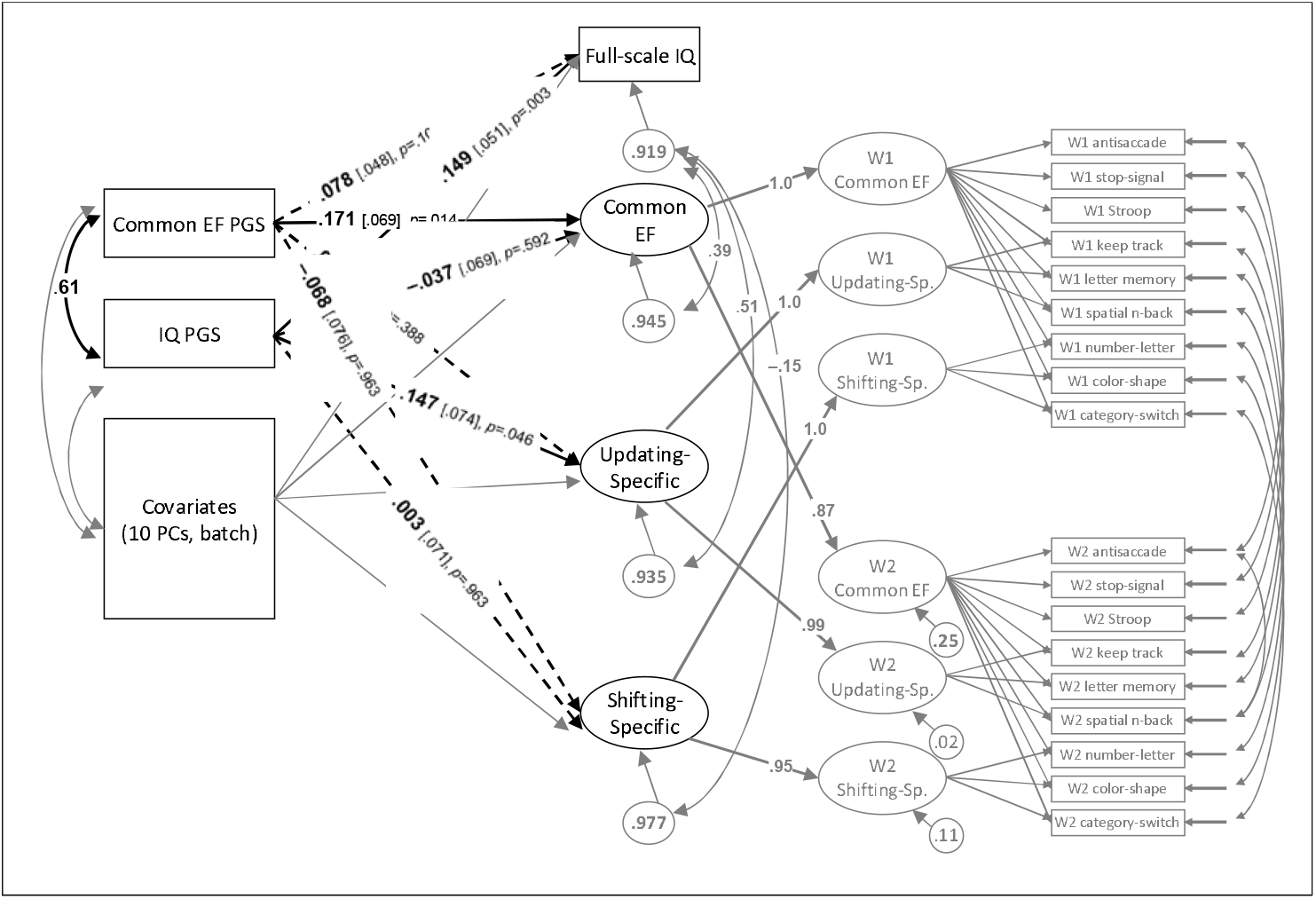
Analysis model of polygenic scores (PGSs) predicting executive functioning (EF) latent variables and full-scale intelligence scores (IQ) in Colorado twin data, χ^2^(350) = 473.85, *p* < .001, CFI = .962, RMSEA = .020. Paths of primary interest are shown in black with thicker lines. Solid lines and boldface type indicate *p* < .05; dashed lines indicate *p* > .05. Analyses were limited to twins with European ancestry based on the first three principal components (*N*=916 with genetic data). The three EF latent variables were based on 9 laboratory tasks at Wave 1 (W1; Longitudinal Twin Study [LTS] age 17 *n*=571, Community Twin Sample [CTS] age 21 *n*=298), and on 9 tasks at Wave 2 (W2; LTS only at age 23, *n*=555). Full-scale IQ was based on 11 Wechsler Adult Intelligence Scale subtests in the LTS (age 16, *n*=584), and 4 Wechsler Abbreviated Scale of Intelligence subtests in the CTS (age 21, *n*=297). Age, sex, and age*sex were regressed out of each measure within each sample and wave prior to analysis.

Controlling for its shared variance with the IQ PGS (*r*=.607, se=.027), the cEF PGS predicted the cEF latent variable in this twin sample (standardized β=.171, *p*=.014, partial *r*=.136), but not the Updating-specific and Shifting-specific factors or full-scale IQ (β= −.068 to .078, *p*>.101, partial *r*s= −.053 to .050). Conversely, controlling for its shared variance with the cEF PGS, the IQ PGS predicted full-scale IQ (β=.149, *p*=.003, partial *r*=.121) as well as the Updating-specific latent variable (β=.147, *p*=.046, partial *r*=. 119), but not the cEF or Shifting-specific latent variables (β= −.037 to .003, *p*>.591, partial *r*s= −.030 to .007). The association of the IQ PGS with working memory updating-specific variance is consistent with prior adult twin studies showing that IQ is genetically related to working-memory-specific latent variables over and above its association with cEF^8,13^. These results are consistent with the hypothesis that the cEF and IQ factors in UKB are tapping similar constructs as those assessed in these carefully phenotyped young adult population samples.

### Genetic Separability of cEF and IQ is Key for Psychiatric Dysfunction

We used LD Score regression^27^ with GWAS summary statistics from previously published studies to estimate the genetic correlation between cEF and other major behavioral and neurological phenotypes (supplemental Table S12), many of which have been associated with EF phenotypically and/or genetically in the literature. cEF was significantly negatively genetically correlated (with Bonferroni correction α=.0011 for 46 traits) with all psychiatric disorders except autism (Figure 4A). 95% confidence intervals of cEF and IQ *r*G did not overlap for five of nine psychiatric traits, but did overlap for neuropsychiatric symptoms, personality, sleep, biometric traits, and most substance use measures.

**Figure 4.**
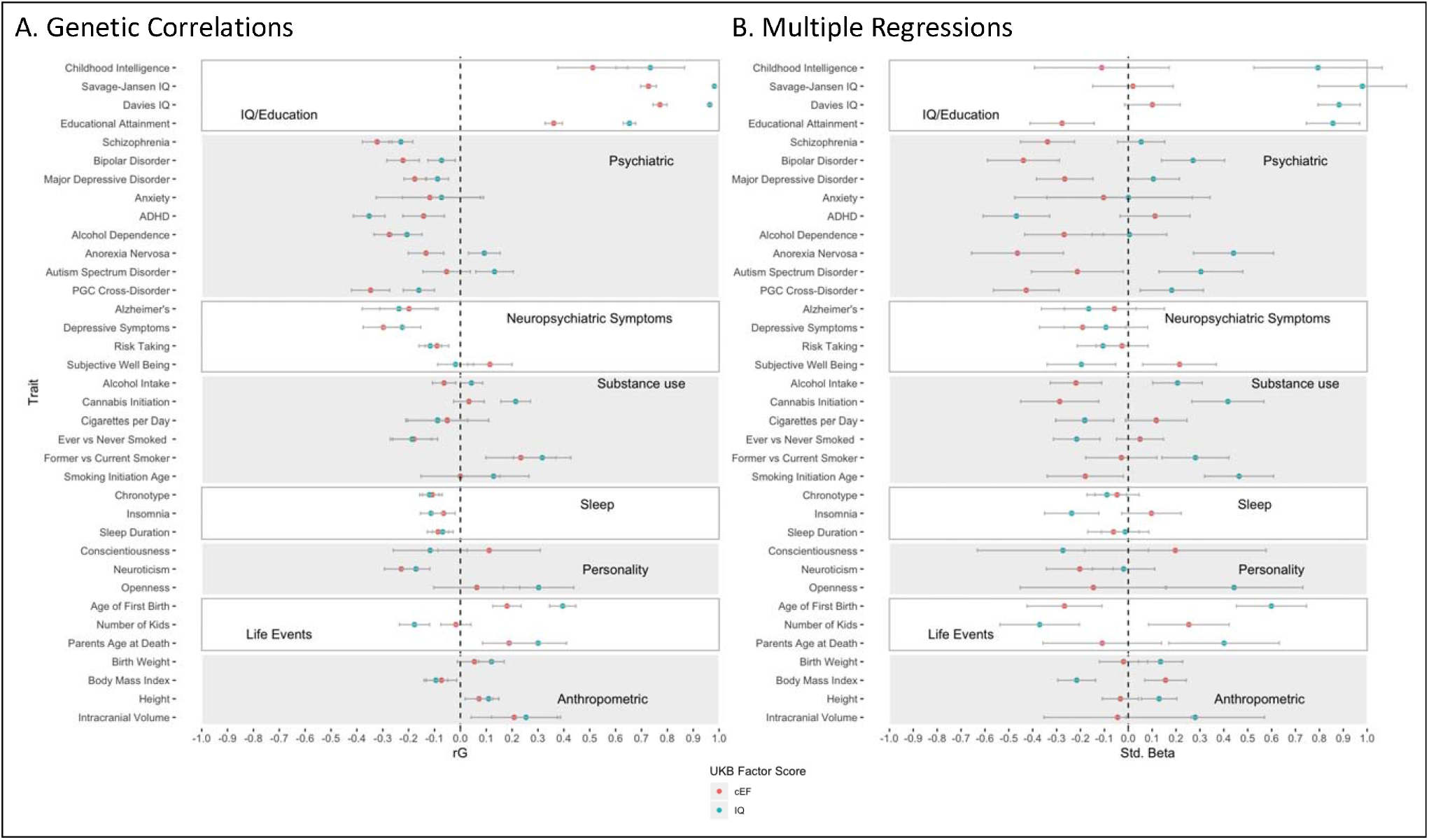
Genetic associations of Common Executive Functioning (cEF) and Intelligence (IQ) factor scores in the UK Biobank (UKB) with psychiatric, behavioral, and health traits in independent samples from LD Hub. (A) genetic correlations, estimated with LD score regression; (B) standardized partial regression coefficients from genomic structural equation models for cEF controlling for the genetics of IQ and Speed, and for IQ controlling for the genetics of cEF and Speed. Bars indicate 95% confidence intervals.

In multiple regressions framework using GenomicSEM^28^ (Figure 4B, Table S13), we estimated significant negative cEF genetic effects on seven of nine psychiatric disorders after controlling for speed and IQ. After controlling for speed and cEF, IQ had a significant negative association only with ADHD and had positive associations with Anorexia Nervosa, Autism spectrum disorder, bipolar disorder, PGC cross-disorder, and (marginally) major depressive disorder. Together, these results suggest that the genes specific to cEF and those specific to IQ have very different influences on the pathogenesis of psychiatric traits.

To formally test the hypothesis that cEF genetic effects are more related to common psychiatric disorders than IQ genetic effects, we estimated a GenomicSEM^28^ model in which UKB cEF, IQ, and Speed predicted a *p* factor estimated from the genetic correlations across psychiatric disorders (Figure 5A; note that GenomicSEM does not provide standard errors for the fully standardized estimates, but we computed the *p*-values based on the STD_Genotype estimates/se in the GenomicSEM outputs). cEF was negatively associated with *p*, controlling for IQ and Speed (fully standardized β= −.50, *p*=1.8e-11), but IQ was no longer negatively associated with *p* (now positive), after controlling for Speed and cEF (β=.12, *p*=.052).

**Figure 5.**
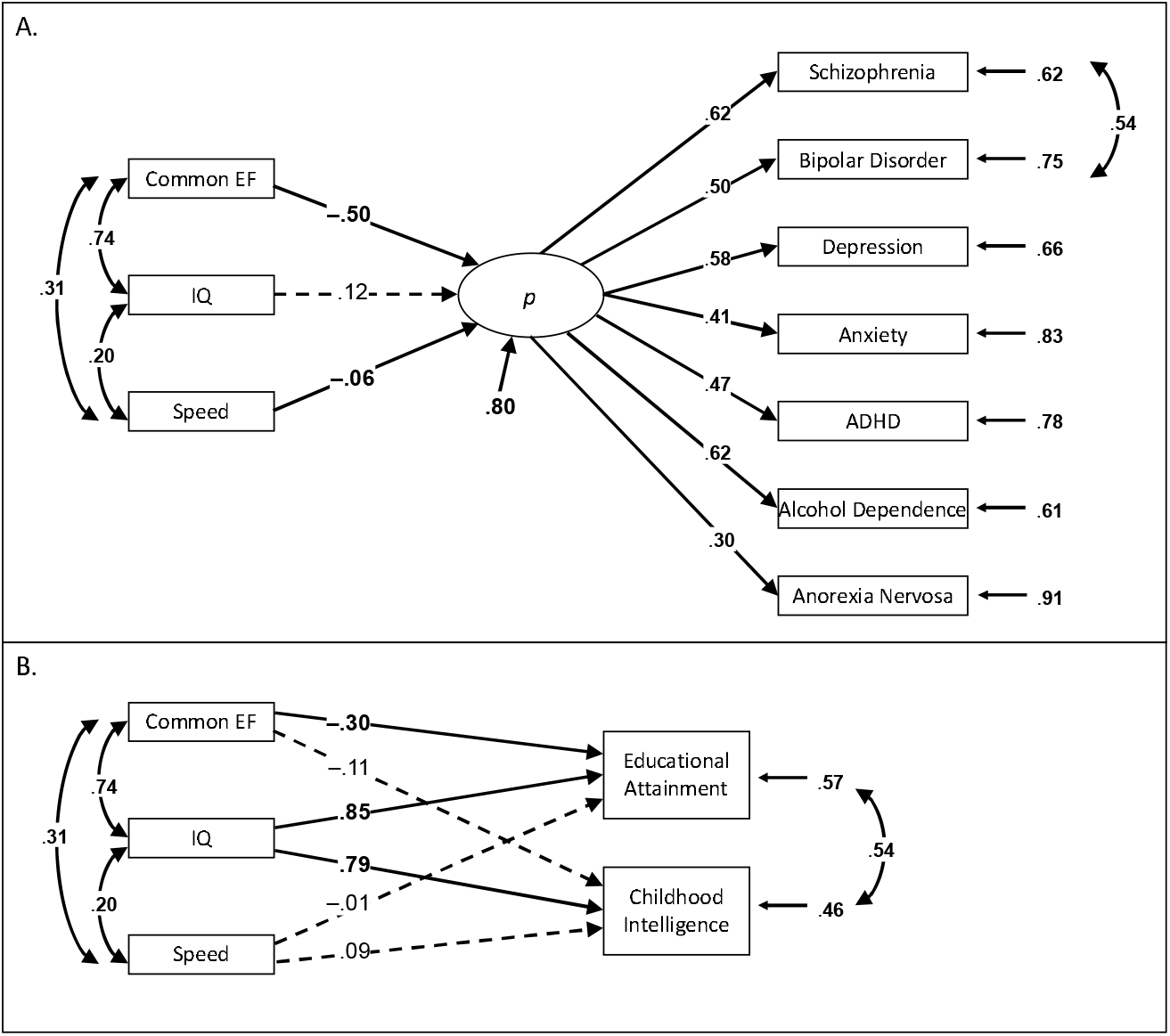
Genomic structural equation models of standardized Common Executive Functioning (cEF), Intelligence (IQ), and Speed factor scores predicting a general psychopathology (*p*) factor (Panel A) and IQ-related traits (Panel B). The fit for the model in panel A was χ^2^(31)= 510.96, *p*<.001, CFI=.901; the model in panel B was saturated, so fit perfectly. Ellipses indicate latent variables; rectangles indicate observed variables. Numbers on single-headed arrows are fully standardized factor loadings or regression coefficients, numbers on curved double-headed arrows are correlations, and numbers at the end of arrows are residual variances. Boldface type and solid lines indicate *p* < .05; dashed lines indicate *p* > .05.

Figure 5B highlights results for traits that show the opposite pattern (also shown in Figure 4): When estimating a GenomicSEM of cEF, IQ, and Speed to predict educational attainment and childhood IQ, IQ was significantly positively related to both, controlling for cEF and Speed (βs= .79 to .85, *p*<7.4e-9); there was a weaker (educational attainment β= −.30, *p*=1.0e-4) or a null (childhood IQ β= −.11, *p*=.438) effect of cEF (Figure 5B and supplemental Table S13). Interestingly, the cEF genetic association with educational attainment changed from positive to significantly negative after controlling for IQ. We discuss this result in the *Discussion*.

Given the relationships between cEF and psychiatric disorders with and without controlling for IQ, we used latent causal variable analysis^29^, a type of genetic causality analysis that uses whole-genome summary statistics, to estimate the genetic causality proportion (gcp) of cEF with schizophrenia, bipolar disorder, MDD, eating disorders, neuroticism, and well-being; these measures were disorders used as indicators in the *p*-factor model (Figure 5), or more general predictors of psychiatric risk that also showed relationships with cEF controlling for IQ in Figure 4B (see supplemental Table S14 for results with the full cEF summary statistics and with the cEF mtCOJO statistics controlling for IQ). We found no evidence for cEF causally influencing any psychiatric disorders, psychiatric dimensions, or other cognitive abilities (all gcp<.42, *p*>.078, with the exception that cEF conditioned on IQ did show a significant influence on Speed, gcp=.18, *p*=.016). These results suggest that the association between cEF and psychiatric disorders may not reflect a simple directional causal mechanism; cEF and IQ are influenced by pleiotropic genes.

### Genetic Associations with cEF Implicate GABAergic and Synaptic Molecular Pathways

We used aggregated gene and gene-set analysis in Multi-marker Analysis of GenoMic Annotation (MAGMA) to understand the GWAS associations. We identified 319 genes significantly associated with cEF in the full sample (Bonferroni α= 0.05/18597=2.689e-6), 21 of which were consistent across the dense and sparse subsamples. The strongest association again was *EXOC4* (supplemental Figures S4-S5, supplemental Table S15).

Using gene-set analyses of this gene list, we found 12 associated gene sets (post-Bonferroni correction), all of which could be summarized under three broad pathways: potassium channel activity, synaptic structure, or GABA receptor activity (Figure 6A, supplemental Table S16). Suggestive associations of additional pathways (corrected *p*<0.1), also implicated synaptic, potassium channel, and ionotropic pathways. To account for some genes appearing in multiple associated pathways, we conducted a conditional gene-set analysis accounting for overlap in genes amongst the top pathways^30^ (excluding the Gene Ontology^31,32^ [GO] terms “synapse,” “GABA_A_ gene,” and “voltage-gated potassium channel” pathways due to multicollinearity). Results indicated that the GO “GABA receptor complex” and “regulation of synapse structure or activity” pathways were associated with cEF over and above other discovered pathways.

**Figure 6.**
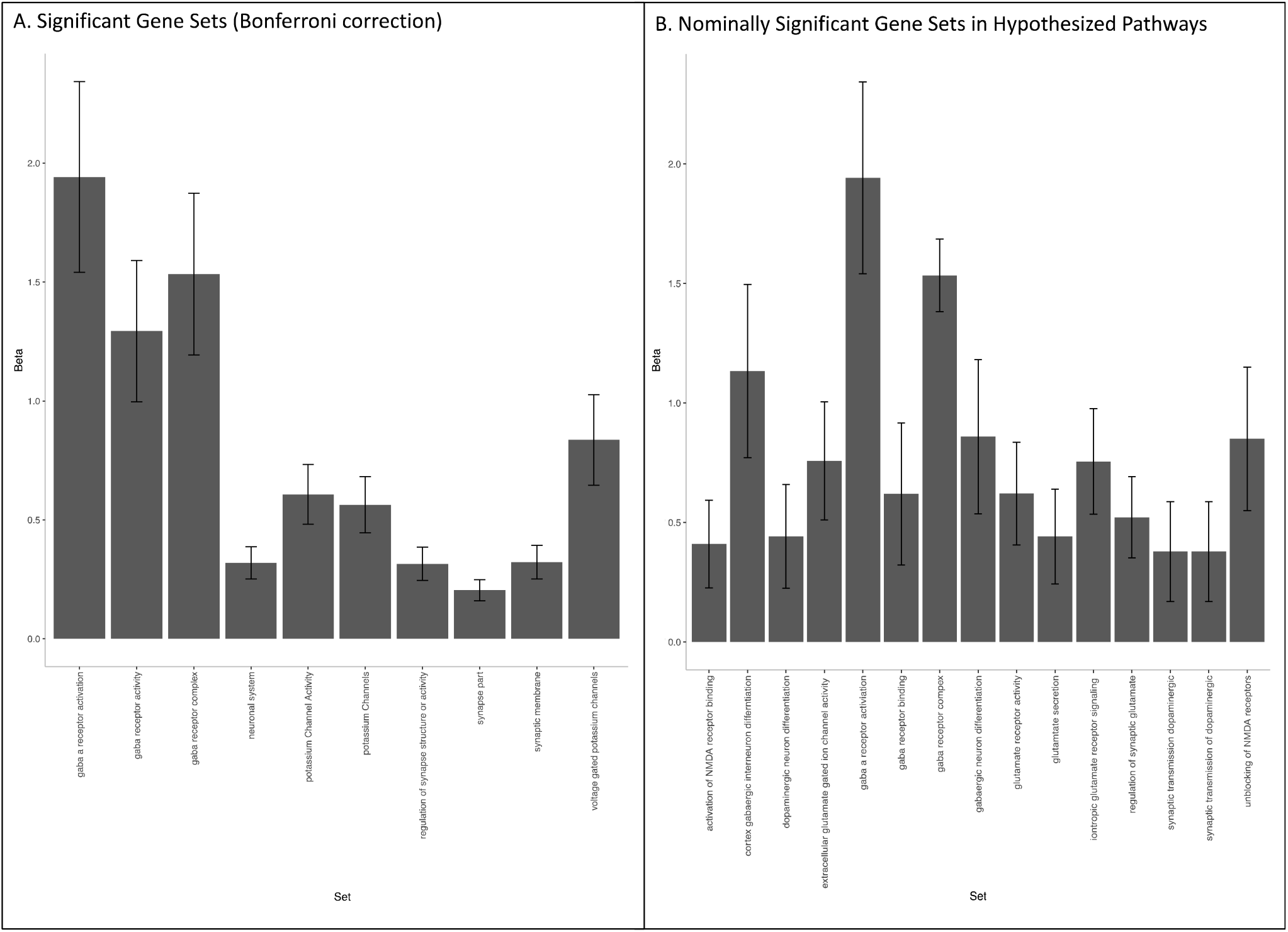
Associated gene-set categories from MAGMA gene-set analysis. Signal GO term and curated gene set enrichment for SNPs influencing Common Executive Functioning (cEF) as the MAGMA gene enrichment beta and standard error. (A) Gene-sets significantly associated post Bonferroni correction for 10,651 tests (α = 4.7E-06); (B) Gene-sets in hypothesized pathways that were nominally significant. VG = voltage-gated. Bars indicate standard errors.

We confirmed that GABAergic function was implicated via cell-type specific gene expression in three human brain tissues: specifically, GABA2 cells in the hippocampus and GABAergic neurons in the prefrontal cortex (specific to 26 weeks of gestation). Hybrid and neurons cells in the whole human cortex (across age) were also significantly enriched for overlap with genes implicated by our GWAS of cEF (supplemental Figure S5).

We expanded our exploration to possible products of genes via a Transcriptome-Wide Analysis (TWAS) using PrediXcan^33^. TWAS uses (cis-)eQTL data to impute gene transcriptomes from available eQTL and GWAS data. Finally, we compared the transcriptomic profile we imputed to the Library of Integrated Cell lines (LINCs) to see if any pertubagens could cause the pattern associated with better cEF. Results are available in supplemental Tables S17-S18 and described in the *Supplemental Results*.

#### Transcriptional differences between cEF and IQ

To assess the distinction between cEF and IQ, we characterized their different expression impacts using overlap with eQTL data, identifying 334,554 cis-eQTLs from the Genotype-Tissue Expression (GTEx)^34^ and the Brain eQTL Almanac (BRAINEAC)^35^ samples. Notably, GRIN2A (the second strongest gene-level association with cEF) expression is altered by ketamine, which has been shown to impair EF performance^19^. We therefore searched for GRIN2A receptor eQTLs within brain regions, finding 10 overlapping cEF associations in the basal ganglia (rs74729488, rs72772387, rs117583711, rs12932206l, rs12597701, rs1213323, rs720717, rs9934226, rs4780790, rs28550823).

The cEF-specific GWAS from mt-COJO showed significant association with only one genetic pathway based on a MAGMA gene-set analysis, Ikeda mir30 targets, which is a pathway involved in the regulation of calcium signaling (*b*=1.008, p_bon_=0.020). cEF was still marginally related to GABA receptor pathways, with a similar effect size as cEF without conditioning on IQ (*b*=1.17, p_bon_=0.079).

### Genetic Associations with cEF Do Not Strongly Implicate Dopaminergic Pathways or Replicate Candidate Genes

#### Test of hypothesized/popular pathways

We hypothesized *a priori* that genes in dopaminergic, glutaminergic, and GABA pathways would be enriched, and it is possible that our multiple comparison threshold was too conservative. We extracted the effects of these pathways that were above nominal significance but did not meet Bonferroni significance. While there were 10 nominally significant pathways from *a priori* categories, the effect sizes and significance values were highest for GABA (Figure 6B). For glutaminergic pathways, the strongest association was NMDA receptor activation, which is previously supported^19^. Finally, dopaminergic genes showed the weakest evidence for association among pre-hypothesized pathways: Only two pathways were nominally associated with cEF, and both pathways showed weaker effect sizes than GABA and glutamate.

#### Candidate gene analysis

Similar to other recent studies on schizophrenia and major depression^36–38^, we found little evidence that the most popular candidate gene polymorphisms (those reviewed by Barnes et al.^17^) were related to cEF at levels above chance. COMT val/met (rs4680), the most-studied candidate gene polymorphism for cEF, was not significant at the genome-wide level (β= −.002, *p*=.021). Previously studied polymorphisms of DRD2 (rs1079596: β=0.010, *p*=1.3e-10; rs2075654: β=0.010, *p*=1.4e-100) were genome-wide significant; however, these effects sizes are much smaller than previously reported^39^. No other associations were even nominally significant. We also used MAGMA to derive combined *p*-values from GWAS summary statistics to determine the degree of association of historical cEF candidate genes themselves as opposed to the most-studied specific polymorphisms within them^17^. Only DRD2 was associated with cEF (*p*=1.15E-12, all gene-wise summary statistics available in supplemental Table S15). Dopaminergic genes other than DRD2 were not significant, suggesting that the weak dopaminergic signal in our pathway analysis above is likely driven by DRD2.

## Discussion

We conducted the largest GWAS of EF to date, using a cEF factor score in the UKB that minimized the task impurity problem and incorporated existing knowledge of the factor structure of EFs^1^. Our results suggest that genetic influences on cEF involve variation within fast ionotropic and synaptic pathways, in particular GABAergic pathways, rather than the commonly studied metabotropic and neuronal pathways. We demonstrated cEF’s genetic overlap with IQ, but found important differences between them as shown through differential associations with education and psychiatric disorders.

In line with existing twin literature^8,13^, this study supports the importance of cEF as a cognitive dimension that is partially genetically related to IQ and Speed in adulthood. Although there was a high genetic correlation between cEF and IQ (*r*G=.743), this correlation was significantly lower than 1, indicating some specific variance. This separability has important implications for understanding cognitive aspects of psychopathology. Controlling for IQ and Speed, cEF remained significantly negatively genetically associated with a genetic *p* factor, accounting for one-quarter of common genetic influences across psychiatric disorders. This stronger association of cEF with *p* cannot be explained as arising from the possibility that cEF may be a more reliable or valid measure of general cognitive ability than the IQ factor score, because the opposite pattern was observed for education and childhood IQ. Specifically, controlling for their genetic overlap, IQ remained strongly positively associated with these measures, while cEF did not. Thus, we observed discriminant validity of the cEF and IQ factor scores at the genetic level, consistent with the phenotypic literature^1^.

Interestingly, when controlling for covariance with IQ, genetic influences on cEF became significantly negatively related to educational attainment, although the genetic correlation without controlling for IQ was positive. We do not know of prior phenotypic studies reporting this suppression effect, nor do we know if it would even be evident at the phenotypic level. It may reflect the fact that cEF shows negative genetic correlations with several disorders that are actually positively genetically correlated with both IQ and educational attainment, such as anorexia nervosa, autism spectrum disorder, and bipolar disorder^25^. Thus, it may be that the genetic variance unique to lower cEF reflects some of this genetic risk for these disorders that is positively associated with education, leading to a negative partial genetic correlation with higher cEF.

Multiple lines of evidence suggested the importance of GABA to cEF variation, with synaptic and ionotropic pathways more strongly associated with cEF than the traditionally studied metabotropic and dopaminergic pathways. GABAergic pathways were the most associated molecular pathways tested and influenced cEF above and beyond other neurotransmitter pathways. Cell-type-specific analyses also implicated GABA in both the cortex and hippocampus. Tests of specificity when accounting for IQ showed that GABA remained marginally related to cEF, with a larger effect size than most other enrichment categories. Together, our findings strongly implicate a key role of fast-synaptic communication mechanisms underlying the inheritance of cEF, rather than the slow neuromodulatory processes that are often hypothesized in the literature.

We found little evidence that dopaminergic processes genetically relate to individual differences in cEF, outside the popular DRD2 gene. Importantly, almost all monoamine (dopamine and serotonin) candidate genes that are currently used in the neurocognitive and imaging literature^17^ were not associated with cEF, despite very high power to detect previously reported associations. These results suggest that continued focus on traditional candidate genes outside of DRD2 in cEF research is likely to be fruitless, and that future research should instead be directed toward its relationship with GABAergic (and perhaps glutaminergic) processes.

It is likely these fast-synaptic processes influencing cEF could further elucidate mechanisms of psychiatric disorders, as cEF was genetically correlated with nearly all psychiatric disorders. We found novel genetic associations with schizophrenia, bipolar disorder, alcohol dependence, and anorexia nervosa, and replicated a genetic association of cEF with depression^40^. These results are in line with past literature, suggesting cEF is broadly genetically associated with psychopathology^2^.

These results should be interpreted in context of a few limitations. First, the cognitive battery in the UKB study was not designed to tap cEF. This battery contained one classic neuropsychological EF task, the trail making task; the other cognitive measures were not tasks that are commonly used to assess EFs. However, as described in the online methods, those measures had EF components that can be extracted through our structural modeling approach. We reasoned that a common factor extracting shared variance across these tasks and the trail making task would be closely related to the Common EF factors examined in smaller studies^9,14–16^, two of which also used the trail making task^9,14^. Results of our PRS analyses support this reasoning: We validated our cEF factor and its separability from IQ using PGSs in a sample deeply phenotyped for multiple EF latent variables and full-scale IQ.

Second, almost all bioinformatic follow-up depended on tissue-based analysis from the GTEx sample (with GABAergic replications from single-cell RNA seq datasets). While this sample is the richest source of eQTL data to date, a lack of generalizability from this population would affect our results as well as our interpretations. Third, we used tissue-specific expression with Bonferroni correction thresholds to minimize false positives, but this does not mean we can draw strong conclusions about which tissues are implicated above and beyond one another.

Finally, because the UKB is overwhelmingly European ancestry, we restricted our analysis to European samples to avoid confounds due to population stratification. Although it is possible and perhaps likely that the molecular underpinnings of cEF generalize to non-European populations, further work is needed to replicate these observations in diverse populations of sufficient sizes and similar phenotypes.

## Conclusion

cEF is heritable and highly polygenic, with clear indication for a role of synaptic, GABAergic, and ionotropic pathways. cEF is genetically related to, but separable from, IQ, and cEF is robustly related to genetic risk for general psychopathology even controlling for its genetic overlap with general IQ and Speed.

## Supporting information

Supplemental Results and Figures

## Data Access

Summary statistics for all GWAS (cEF, IQ, RT, and all indicators) will be available upon publication of this work through the FriedmanLab github or available upon request. All results run on FUMA have been made publicly available through that platform. cEF results can be accessed via FUMA (cEF full sample: https://fuma.ctglab.nl/browse/65, cEF densely phenotyped sample: https://fuma.ctglab.nl/browse/66, cEF sparsely phenotyped sample: https://fuma.ctglab.nl/browse/67). Full results for the IQ and Speed GWAS are also available on FUMA (IQ: https://fuma.ctglab.nl/browse/114, Speed: https://fuma.ctglab.nl/browse/118). All biological results for cEF-specific and IQ-specific GWAS can be downloaded here (cEF-specific: https://fuma.ctglab.nl/browse/116, IQ-specific: https://fuma.ctglab.nl/browse/117).

## Acknowledgements

This research was supported by grants from the National Institutes of Health: MH063207, MH016880, MH001865, MH100141, AG046938, DA042742, and DA046064. We thank Mengzhen Liu, Marie Banich, and Marissa Ehringer for their advice. We also thank Cold Spring Harbor Laboratory for hosting this manuscript on their preprint server, bioRxiv.

## Author Contributions

Alexander S. Hatoum ran the genome-wide association analysis and analyses with summary statistics along with Evan C. Mitchell and Claire L. Morrison, and wrote the manuscript along with Naomi P. Friedman. Naomi P. Friedman sponsored the research, generated the phenotype in the UKB, conducted the PGS analyses along with Chelsie E. Benca-Bachman, and co-wrote the manuscript. Luke M. Evans organized the raw genotype data from the UKB as well as the genotype data in the twin samples. Matthew C. Keller, Luke M. Evans, Andrew E. Reineberg, Max Lam, and Rohan H. C. Palmer provided advice and assistance with analyses and manuscript preparation. All authors contributed to the final written version of the manuscript.

