## Supplemental Results and Figures for "Genome-Wide Association Study of Over 427,000 Individuals Establishes Executive Functioning as a Neurocognitive Basis of Psychiatric Disorders Influenced by GABAergic Processes"

**Supplementary Results and Figures**

***Common Executive Functioning (cEF) and Gene Expression Patterns***

We localized expression patterns to 11 brain regions from the 53 human tissues in the GTeX sample (all except the substantia nigra and spinal cord-c1, supplemental Figure S6). we used PrediXcan^1^ to predict brain transcription patterns of higher cEF from our SNP summary statistics and tissue-specific eQTL expression associations from the GTEx sample's^2^ 11 associated brain tissues. We found 441 brain tissue-specific transcripts (of 4,324 possible) associated with cEF, post-Bonferroni correction (supplemental Table S17, expression pattern across tissues in GTEx in supplemental Figure S7).

We then entered this transcriptional profile in the connectivity Map (cMAP)^3^. After filtering for transcripts found in multiple tissues, 78 connectivity map loci that were also associated with transcriptional changes after exposure to perturbagens in the cMAP. Thirty-three perturbagens mimicked or reversed the cEF transcriptional profile (supplemental Table S18). Of note, 3 of the top 15 substances have previous psychiatric and cognitive applications: nicergoline^4^, an anti-dementia drug that is shown to be effective in a broad array of behavioral and cognitive disorders in old age; nortriptyline^5^, a first-generation tricyclic antidepressant; and chlorpromazine^6^, a typical anti-psychotic that is prescribed to treat severe cases of schizophrenia, bipolar, obsessive-compulsive disorder, and depression.

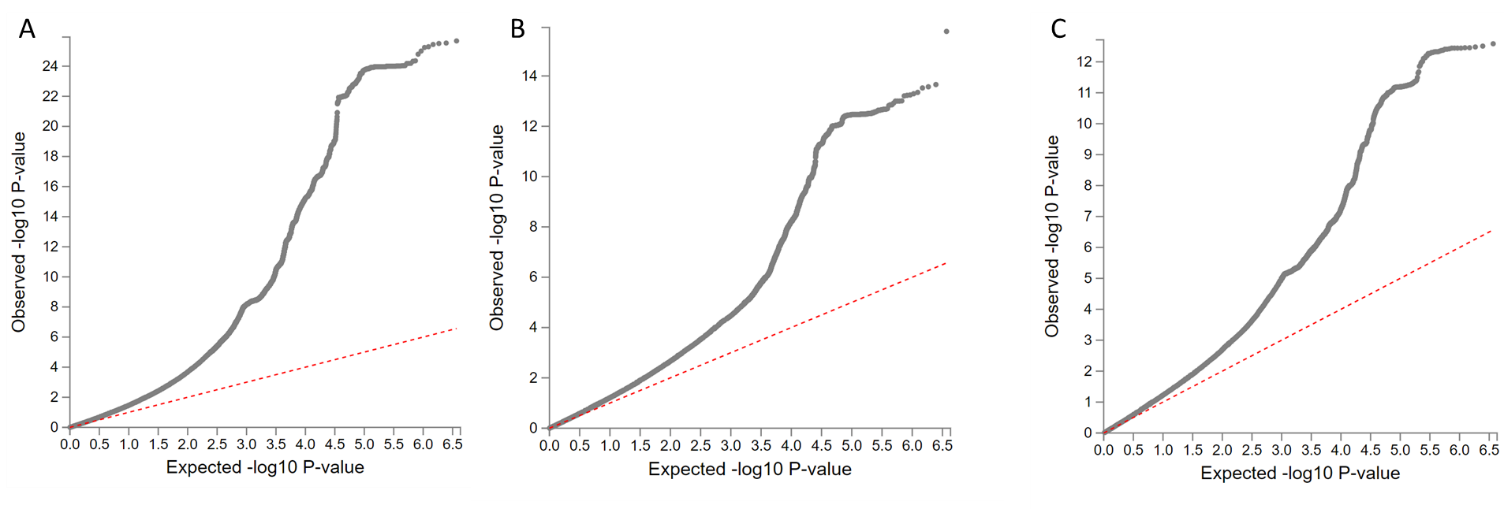

**Supplemental Figure S1. QQ plots of SNP *p*-values for GWAS of Common Executive Functioning (cEF) factor scores in the (A) full sample; (B) deeply phenotyped sample (B); and (C) sparsely phenotyped sample.** Black dots represent the observed *p*-values plotted against the y axis on the –log10 scale, red dots represent the expected *p*-values plotted on the x-axis; *p*-values deviated substantially from expected.

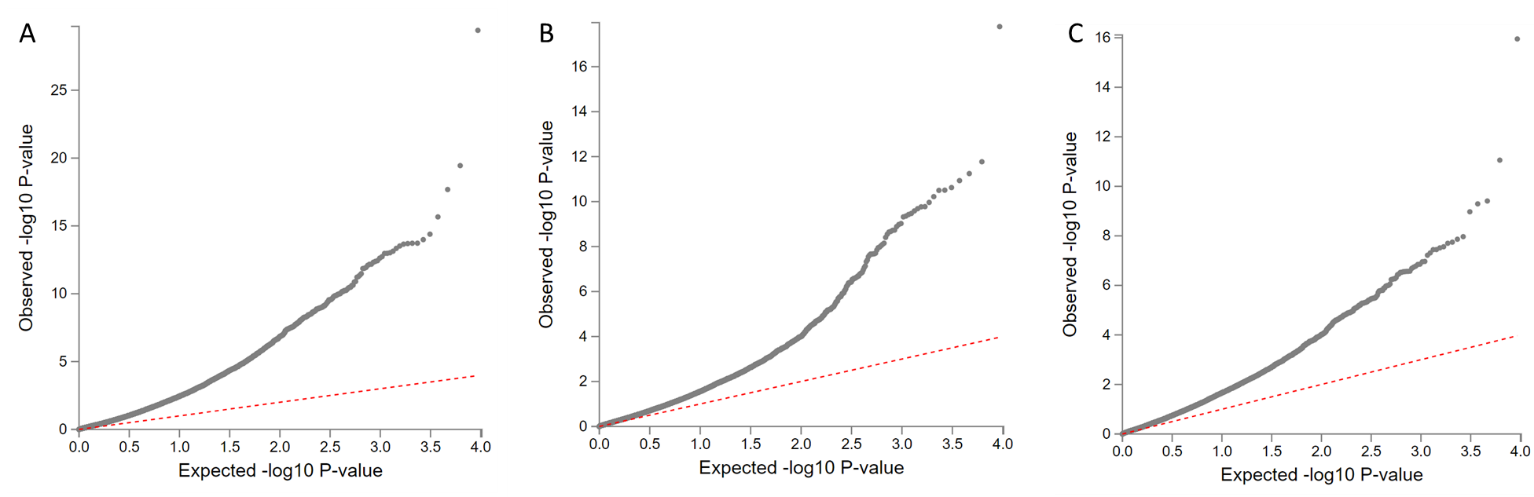

**Supplemental Figure S2. QQ plots of gene-wise *p*-values for Common Executive Functioning (cEF) factor scores in the (A) full sample; (B) deeply phenotyped sample (B); and (C) sparsely phenotyped sample.** Black dots represent the observed *p*-values plotted against the y axis on the –log10 scale, red dots represent the expected *p*-values plotted on the x-axis; *p*-values deviated substantially from expected.

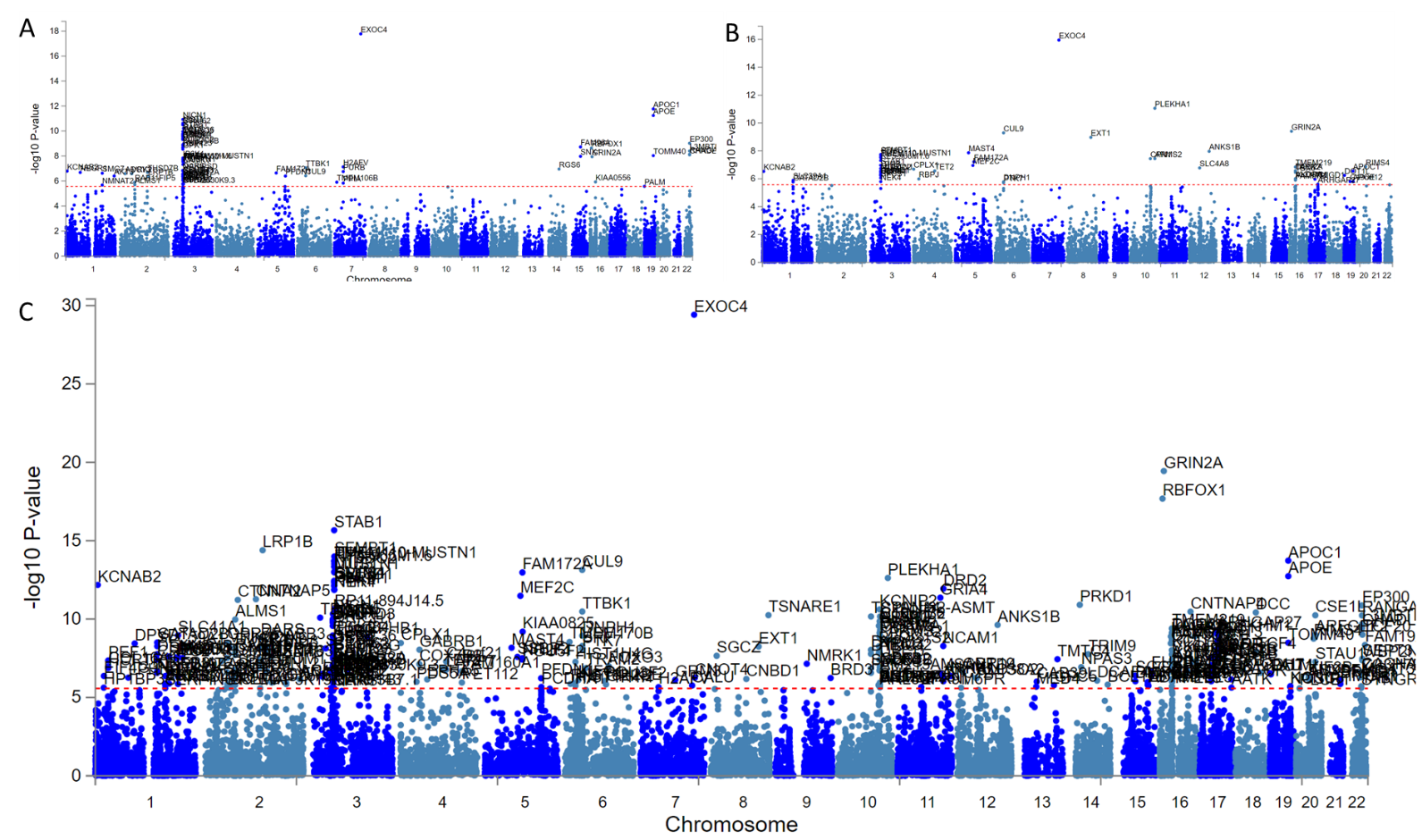
**Supplemental Figure S3. Manhattan plots for three gene-wise association tests of Common Executive Functioning (cEF) factor scores.** The UK Biobank was split into (A) a discovery sample that was densely phenotyped; and (B) a sparsely phenotyped sample. Relatives were pruned from the sparsely phenotyped sample to ensure gene associations were not due to inflation by cryptic relatedness. The genes must have been associated by GWAS Bonferroni significance (*p* = 0.05/18739 = 2.668e-6). (C) Full sample: Both samples were combined, and related individuals were included to increase power. All models were run using Bolt-LMM to account for polygenicity and family structure.

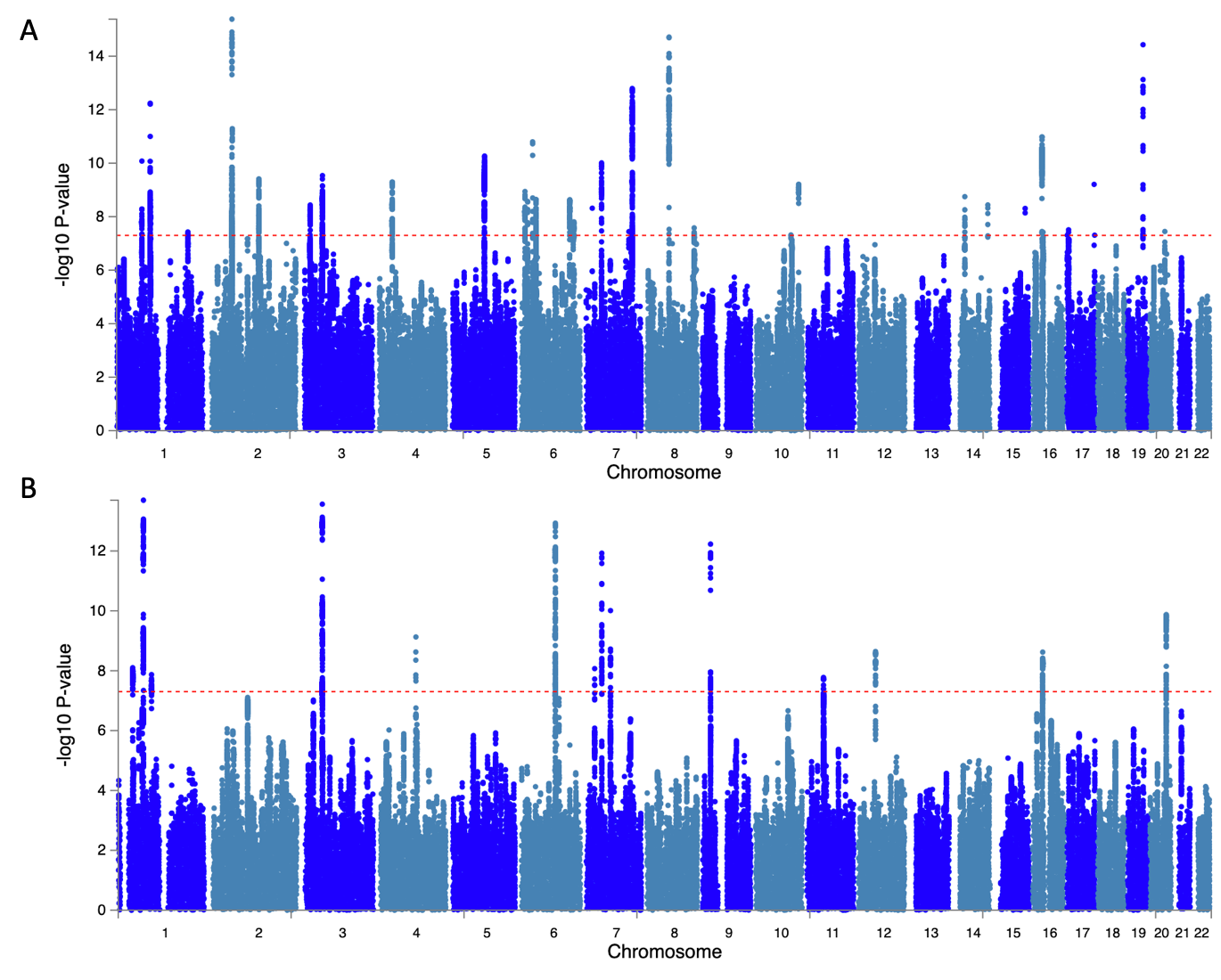
**Supplemental Figure S4**. **Manhattan plots for (A) Common Executive Functioning (cEF)-specific GWAS; and (B) Intelligence (IQ)-specific GWAS,** conducted with mt-COJO. Red dotted line represents genome-wide significance threshold.

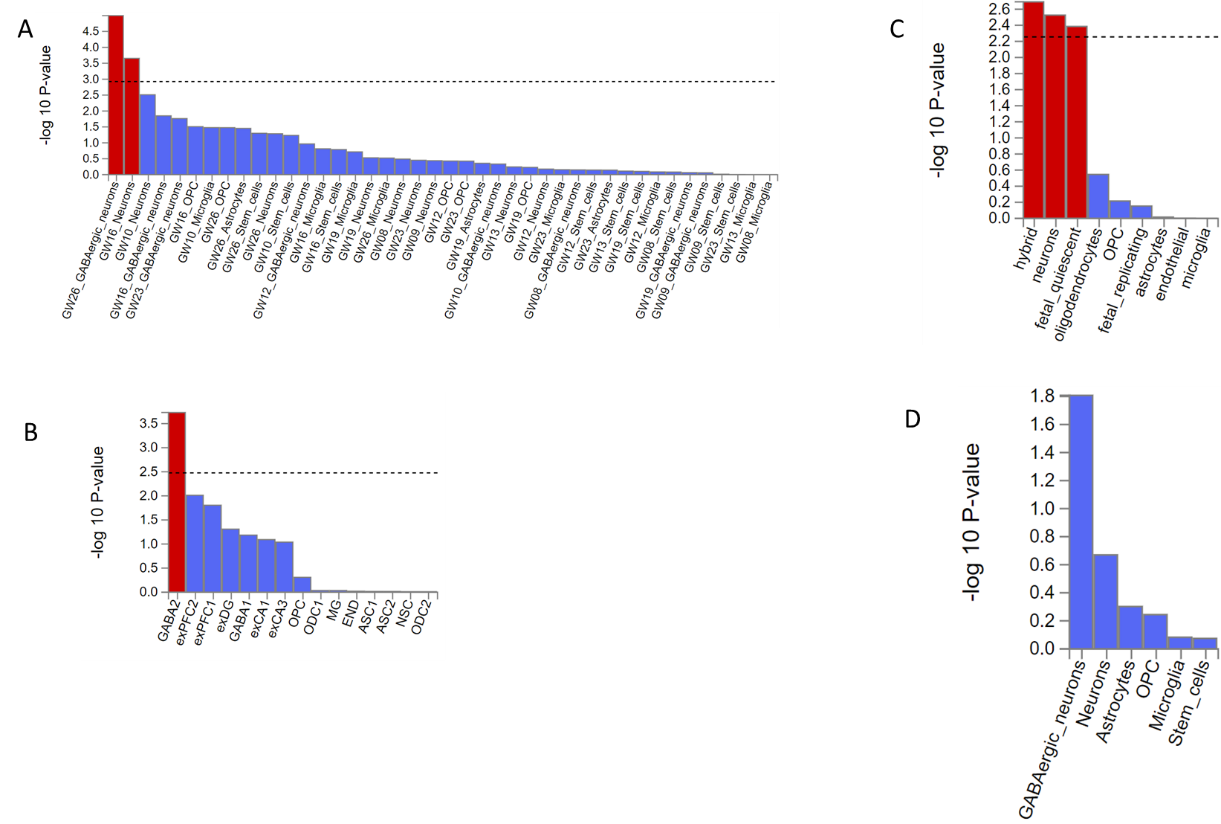

**Supplemental Figure S5. Common Executive Functioning (cEF) RNA-cell type-specific enrichment in human post-mortem brain samples**. To identify cellular mechanisms of cEF SNP associations we used MAGMA gene-set analysis to predict four different post-mortem datasets. (A) Cortex expression patterns across fetal development; (B) Cell-type specific analysis in human post-mortem hippocampal tissue; (C) Combined cortex expression across time (adults and fetal tissue) to predict what cell-types (neuronal/immune) related to cEF; (D) Cell-type-specific analysis in the human post-mortem prefrontal cortex tissue.

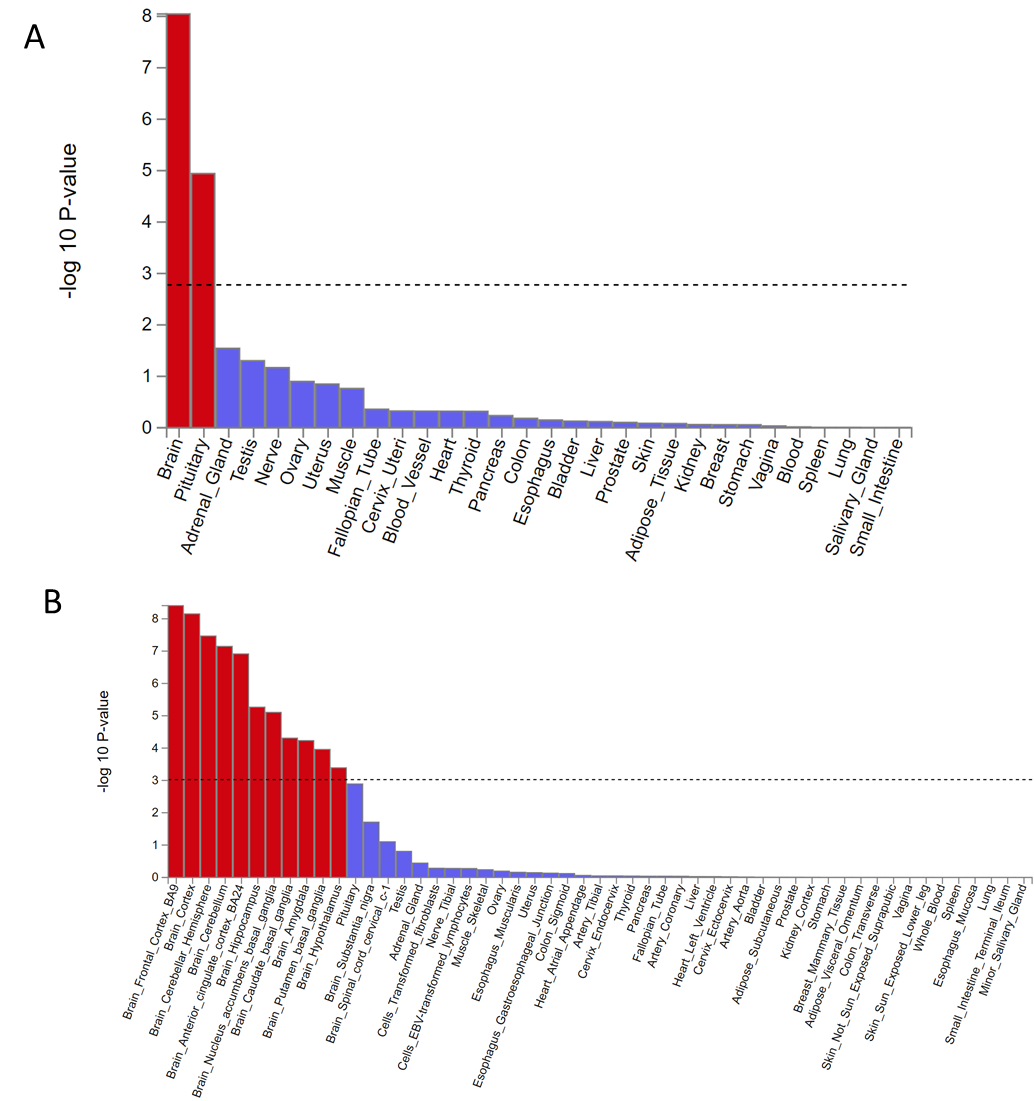

**Supplemental Figure S6. Significant tissue enrichment for Common Executive Functioning (cEF) SNP eQTLs in the GTEx v7 sample.** Enrichment was estimated with MAGMA using a gene-level regression controlling for gene-size and population structure. Line represents Bonferroni significance. (A) Enrichment by broad general tissue types; (B) Enrichment by 53 specific tissue types.

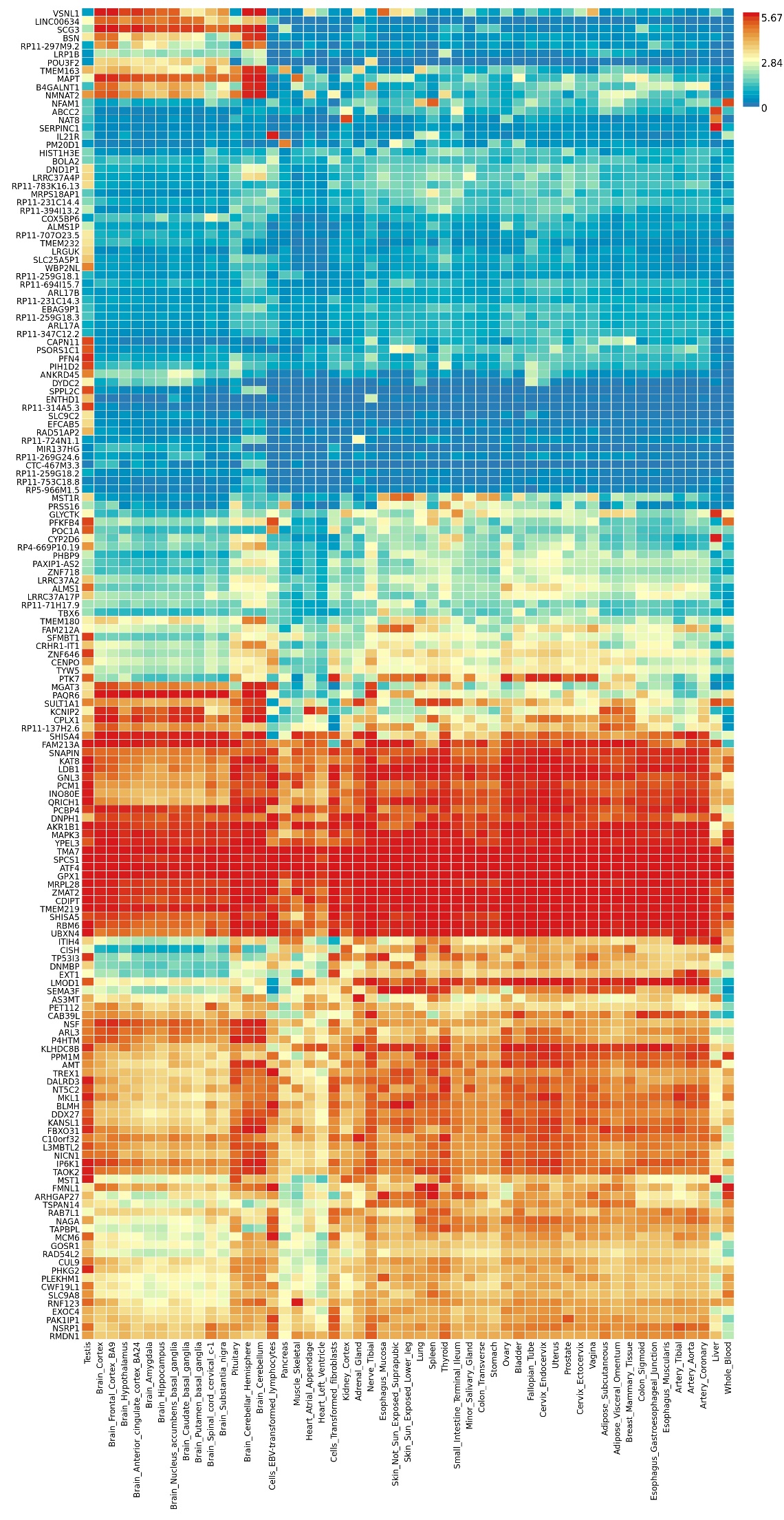

**Figure S7. Transcriptional profile for Common Executive Functioning (cEF) genes as a co-expression heatmap across 53 specific GTEx tissues.** Results are clustered based on tissue and gene expression.

**Supplemental Results References**
