## Supplementary material for "Genome-Wide Association Study of Over 427,000 Individuals Establishes Executive Functioning as a Neurocognitive Basis of Psychiatric Disorders Influenced by GABAergic Processes": Online Methods

**Participants**

Participants were 501,826 individuals in the UK Biobank (UKB) study^1,2^ who had completed at least one cognitive assessment at the time that the data were released to us. The UKB includes a total of 502,544 participants (54.4% female) aged 56.5 years (SD = 8.1, range = 37-73) at recruitment. Cognitive data were collected at up to four time points: an initial assessment visit (2006-2010) during which participants completed cognitive function tests on a touchscreen computer, a repeat assessment (2012-2013), an imaging visit (2014+), and a cognitive online follow-up (2014+). Sample sizes differed for each visit and task described below, as detailed in Table 1. Notably, the sample had dense cognitive assessment at the online follow-up (93,024 individuals overall).

We restricted genetic analyses to 427,037 individuals of European ancestry as determined by principle components (PC) analysis (mean age = 56.849, SD = 8.009, 54% female, 46% male) whose genotypes were imputed to the Haplotype Reference Consortium^3^, 1000 Genomes, and UK10K reference panels by the UKB^2^. Subjects were genotyped on a UK BiLEVE array or the UKBiobank axiom array^2^. After removing individuals with mismatched self-reported and genetic sex, we filtered imputed single nucleotide polymorphisms (SNPs) using a Hardy-Weinberg equilibrium threshold of *p* < 1×10^6^, variant missingness > 0.05, imputation quality score (INFO) > 0.95, and minor allele frequency (MAF) > 0.01, retaining 7,391,068 SNPs.

To guarantee consistent SNP effects across missingness patterns and replicate SNP effects, we had three phases of genome-wide association study (GWAS) analysis. First, we conducted our GWAS in the full sample (*n* = 427,037, mean age = 56.849, SD = 8.009, 54% female). Then, to evaluate consistency across subsamples, we divided the sample into a densely assessed subsample (*n* = 93,024, mean age = 56.065, SD = 7.657, 55% female) and a sparsely assessed subsample. To be in the densely assessed sample, individuals must have completed at least the trail making task, which is a classic neuropsychological EF task that has been used to tap cEF factors in prior studies^6,8^. The sparsely phenotyped sample consisted of the remaining individuals who completed at least one neurocognitive task and were unrelated to people in the densely phenotyped sample. We pruned for related individuals in the sparsely assessed sample (using PLINK's "greedy" algorithm^4^) so SNP discovery and the genetic correlation between subsamples would not be biased by related individuals (final sparsely assessed subsample *n* = 256,135, mean age = 56.996, SD = 8.050, 54% female). We found consistency and independent SNP replication between the subsamples, so we conducted all follow-up analysis on the full sample to maximize power.

**Measures**

**Executive function (EF) measures.**

The cognitive battery in the UKB contains one classic neuropsychological EF task, the trail making task. The other cognitive measures were not tasks that are commonly used to assess particular EFs, but a number of them have EF components that can be extracted through our structural modeling approach. These tasks were symbol-digit substitution, digit span, prospective memory, and pairs memory. We reasoned that a common factor extracting shared variance across these tasks and the trail making task would be closely related to the Common EF factors examined in smaller studies^5–8^, two of which also used the trail making task^6,8^.

**Trail making (online).** Participants clicked the computer mouse to join sets of circles "as quickly and accurately" as possible. The first set (numeric) consisted of 25 circles enclosing the numbers 1-25, which participants joined in ascending order. The second set (alphanumeric) consisted of 25 circles enclosing numbers and letters (1-13 and A-L), which participants joined in alternating order (1, A, 2, B, 3, C, etc.). Each set was preceded by a practice set of 8 circles. The total time in seconds taken to correctly complete each set was recorded, starting when the participant clicked the first item. Only correct answers were accepted; incorrect answers were recorded but not analyzed here, because the time to commit errors was included already in the total time. The alphanumeric set is a classic neuropsychological EF measure because it requires participants to avoid the prepotent tendency to join stimuli in ascending order; rather, participants must switch between two sets of stimuli and maintain and update information about the current position within each set. The numeric set is used as a control condition to assess variation due to basic processing and motor speed. The dependent measure (DM) was the log-transformed time to complete the alphanumeric set (field 20157) residualized on the log-transformed time to complete the numeric set (field 20156).

**Symbol-digit substitution (online).** Participants saw a grid with 8 symbols above the digits 1-8, presented left to right, at the top of the screen. Underneath that, they saw the symbols re-arranged, and had to place the numbers 1-8 underneath them using the keypad, "as quickly and accurately" as possible. After a practice set, participants had 1 minute to complete as many grids as possible. The DM was the number of symbol-digit matches made correctly (field 20159); data from individuals who "abandoned" (field 20245) the test were treated as missing. Although this test is often used as a processing speed measure, its requirements to avoid the prepotent tendency to enter numbers in order and to update which symbol is paired with which number across grids are somewhat executive in nature. Supporting this conceptualization, trail making performance, a classic measure of EF that controls for basic speed, correlated more strongly with symbol-digit substitution (*r* = –.34) than with the simple reaction time (RT) measures we describe below (*r* = .16 to .18 for the 3 RT assessments). Thus, we included this test in both the EF (standardized loading = .606) and RT factor score models (standardized loading = .383).

**Prospective memory (initial visit, repeat assessment, imaging visit).** Part 1: Before any other cognitive tests, participants saw the following text, "At the end of the games we will show you four coloured shapes and ask you to touch the Blue Square. However, to test your memory, we want you to actually touch the Orange Circle instead." Part 2: After they completed the other cognitive tests, they saw the following text, "That's the last game. Just one more thing left to do…” After they pressed "Next", they saw a screen with four shapes (blue square, pink star, gray cross, and orange circle), along with the instruction, "Please touch the Blue Square then touch the 'Next' button." If they pressed "Next" without touching a shape, they were prompted to touch a shape. The symbol they touched was surrounded by a yellow box. If they touched any symbol besides the blue square, the test ended, but if they touched the blue square, they received the following instructions, "At the start of the games we asked you to remember to touch a different symbol when this screen appeared. Please try to remember which symbol it was and touch it now." This prompt repeated each time they touched the blue square, until they touched any other symbol, which ended the test. The DM was whether they touched the orange circle on the first try (field 20018). Data from individuals who "abandoned" (field 4287) the test were treated as missing. If they never touched the orange circle or touched it on the second try, it was scored as incorrect. Although this test assesses memory, we judged it to also have an EF component because that memory is for a goal that must be used to override the more salient current instruction to touch the blue square.

**Pairs memory (initial visit, repeat assessment, imaging visit, online).** In the first round of this task, participants had 3s to memorize an array of 6 cards with 3 symbols (i.e., 3 pairs) displayed in a random order. The cards were then shown face-down, and participants had to select the matching pairs. After they touched 2 cards, they were turned over. If they matched, the pair disappeared and the participant selected another pair. If they did not match, they were turned face-down again and the participant touched another pair. This procedure continued until all pairs were correctly identified. After the 6-card round, participants completed a 12-card (6 pairs) round. The DM was the log-transformed number of incorrect matches in the round +1 (field 399 for in-person and 20132 for online), summed across the 6- and 12-card rounds. Data were treated as missing if participants did not match all of the pairs in a round (e.g., if they abandoned the task; field 398 for in-person and 20131 for online), and the DM was only created for participants who had complete data for both rounds. Although this task taps visuospatial memory, it also requires working memory maintenance and updating ability, particularly as incorrect pairs are revealed, so we included it as an EF measure.

**Digit span (initial visit, imaging visit, online).** The digit span test (called "numeric memory" by UKB) required participants to recall numbers with increasing numbers of digits (from 2 to 12). Participants were shown a number for 2s + 500ms*number of digits (e.g., 3s for a 2-digit number). The number disappeared and after 3s, the participant entered the number. After pressing "Next," the entry was removed and the next number was presented 600ms later, or the test was ended. The number length increased by 1 digit with each correct answer, and the test was terminated after 2 successive incorrect answers (if 3 or more digits), or 5 successive incorrect answers if 2 digits. The keyboard was deactivated when the entry screen was not present. Each number was different than the previous number and the previous but one number. The DM was the maximum number of digits remembered correctly (field 4282 for in-person and 20240 for online); data from individuals who "abandoned" the test (coded as a score of –1 and/or "abandoned" in field 4281 for in-person) were treated as missing. The digit span is a classic test of verbal short-term memory, which is not generally considered executive (unless the backward span is used, which requires working memory to re-arrange); typically, complex working memory span tasks that have a simultaneous processing requirement are used to tap working memory EF processes. However, as argued by Unsworth and Engle (2007)^9^, simple and complex working memory tasks seem to measure similar processes (e.g., working memory maintenance, updating, and controlled retrieval) but differ in the extent to which those processes operate. Moreover, as the number of digits exceeds short-term memory span (supraspan), the task becomes more predictive of higher-order cognition. Thus, we included it as an EF measure.

**Other cognitive measures.**

**Intelligence (IQ; initial visit, repeat assessment, imaging visit, online).** In the “fluid intelligence/ reasoning” test, participants had 2 minutes to answer as many questions as possible in a sequence of 13 questions (e.g., "Add the following numbers together: 1 2 3 4 5" and "If Truda's mother's brother is Tim's sister's father, what relation is Truda to Tim?"). The DM was the sum of the correct answers (field 20016 for in-person and 20191 for online). Data from individuals who "abandoned" the online version (field 20242) were treated as missing.

**Speed (initial visit, repeat assessment, imaging visit).** The reaction time (RT) measure (called the Snap game by UKB) was designed to assess simple processing speed. Participants viewed two cards at a time for 2s, and pressed a button on a button box as quickly as possible when they matched. There were 12 pairs of cards, and the first 5 pairs were treated as practice and not included in the score; the pilot phase of recruitment included 20 pairs instead of 12. The DM was the log-transformed mean time to correctly identify matches (field 20023) across the remaining 7 pairs, excluding pairs with RTs < 50ms (anticipatory responses) and >2,000ms (responses that occurred after cards had disappeared). We excluded mean data from participants who never (*n* = 2449) or always (*n* = 261) pressed the button (based on the maximum and minimum values for number of times the button was pressed in field 403, or 10141 for the pilot phase), as this indicated they were not following instructions. We also excluded mean data when the mean was below 300ms (*n* = 42), as we deemed this an unrealistic average time given the overall distribution (mean = 561ms, SD = 117).

**General Procedure**

At the assessment visits, participants completed the touchscreen cognitive tests after reception, consent, a touch-screen questionnaire, and a hearing test. The order was prospective memory part 1; pairs memory; IQ; RT, prospective memory part 2. The digit span (numeric memory) task was phased out of the battery towards the end of recruitment so was only available for a subset of participants. For the online follow-up, the IQ, pairs memory, and digit span tests were implemented as web-based questionnaires to be completed remotely, and the trail making and symbol-digit substitution tests were added. The order was IQ, trail making, symbol-digit substitution, pairs memory, and digit span. The RT test was also included, but the UKB did not release those data because of poor quality.

**Analysis Procedures**

**Factor scores.**

After log-transforming skewed variables, values greater or less than 4 SDs from the mean were replaced with values equal to 4 SDs from the mean. Given the sample size involved, this trimming procedure had little influence on the correlations or model results, but it improved the normality of the distributions (see Table 1), which is an assumption of structural equation modeling. Variables were rescaled to have variances close to 1 to avoid ill-scaled matrices, which can cause model non-convergence, and variables were reversed when appropriate (the DMs for trail making, pairs memory, and RT) so that for all measures, higher scores indicate better performance.

We used Mplus version 8 for the confirmatory factor analyses and extracted factor scores for the Common EF (cEF) factor shown in Figure 1, as well as IQ and Speed factors. These factors were estimated separately, rather than within the same model, to avoid inflating the correlations among the factor scores; factor scores incorporate all correlated variables in the model and factor score indeterminacy can lead to increased correlations among the factors scores, compared to the estimates within the latent variable models from which they are extracted.

The IQ and Speed models used full-information maximum likelihood estimation, which uses all available data and assumes missing data are missing at random. For the cEF model, the prospective memory scores were categorical (pass/fail); we analyzed them with probit link function, which assumes these categories reflected an underlying normal distribution of probability of remembering. These model used means- and variances-adjusted weighted least-squares estimation (WLSMV). Thus, Table 1 presents tetrachoric and biserial correlations with prospective memory performance. For WLSMV, only pairwise deletion is available.

Factor scores were computed for participants with at least one indicator. In the case of individuals who only participated in the first assessment, their factor scores for cEF might be based only on their scores for the pairs memory task, as that was the only task given to the full sample. In the case of individuals with multiple assessments including the online battery, their factor scores would be based on a weighted combination of multiple tasks. The latter are better factor scores because more information is available, and the uncorrelated error variances across tasks cancel each other out (their expected mean is zero); this lower error variance means that the densely phenotyped cEF factor scores should show a higher heritability estimate. Thus, after extracting cEF factor scores from the entire sample, we split them into those obtained from individuals who had completed the trail making test in the online battery and all others as described in the Participants section.

Figure 1A presents the zero-order correlations among the cognitive measures used in the factor models, as well as the factor scores. The confirmatory factor analyses used to obtain the cEF scores is shown in Figure 1B. For the cEF model, all tasks loaded on the cEF factor, and orthogonal task-specific factors were used to account for the fact that some tasks (prospective memory, pairs memory, and digit span) were repeated. We did not conduct GWAS on factor scores for the specific factors, which can be considered to reflect a combination of method variance as well as variance due to processes specific to that paradigm (i.e., uncorrelated with the other tasks in the model). Model fit was good, χ^2^(44) = 1786.53, *p* < .001, CFI = .980, RMSEA = .009. With this sample size (total *n* = 490,588), a large chi-square statistic is expected, but the model fit well by other fit criteria, particularly a CFI > .95 and RMSEA < .06^11^.

We also investigated whether additional factors were needed to capture time-specific effects. When these factors were included, they had non-significant loadings.

Table 2 provides the genetic correlations of the EF tasks with each other. For this table only, we reduced the data to obtain a single score for duplicate tasks as follows: If multiple instances of the same task were available (i.e., pairs memory, digit span), they were standardized within wave, then averaged across waves. As the prospective memory task was categorical, we did not average it across waves; we just used the scores for the first assessment visit. The EF tasks were all genetically correlated with each other (*r*G = 0.3226 to .7126), supporting the use of the latent variable framework.

The IQ and Speed models were primarily based on multiple measures of the same task, so additional factors were not possible. The IQ model (total *n* = 250,544) fit well, χ^2^(2) = 6.83, *p* = .033, CFI = 1.00, RMSEA = .003. The standardized loadings on the IQ factor were .81, .83, .83, and .77 for assessments 1, 2, 3, and online, respectively (all *p* < .05). The Speed model (total *n* = 496,853) also fit well, χ^2^(2) = 0.94, *p* = .625, CFI = 1.00, RMSEA = .000. The standardized loadings on the Speed factor were .74, .81, .76, and .38 for first RT, repeat RT, imaging visit RT, and online symbol-digit substitution assessments, respectively (all *p* < .05).

**Genome-wide association analysis.**

We followed the same procedure for GWAS of the full sample cEF, densely phenotyped sample cEF, sparsely phenotyped sample cEF, Speed, and IQ factor scores. We ran a test of association using a leave-one-chromosome-out Bayesian approximation of a linear mixed effect model using BOLT-LMM, controlling for age, age^2^, sex, first 20 principal components (PCs), batch, and site. BOLT-LMM is a faster and more statistically powerful procedure for running GWAS in large samples (compared to standard software like PLINK and GCTA) and has demonstrated high efficiency with the UKB^12^. This procedure better accounts for stratification/cryptic relatedness and family structure than simply using fixed-effect PCs. The summary statistics in analyses of SNP effects used BOLT’s LMM infinitesimal model *p*-values^13^. Genome-wide results were entered in the Functional Mapping and Annotation (FUMA) / Multi-marker Analysis of GenoMic Annotation (MAGMA)^14^ pipeline^15^, linkage disequilibrium (LD) score regression^16^, and PrediXcan^17^ to characterize the results. Each method is explained below.

Because cEF and IQ correlated latent factors from multiple cognitive tasks, we used multi-trait-based conditional & joint analysis (mtCOJO, within the GCTA-GSMR family of methods)^18^ using GWAS summary data to discover SNP effects that were related to cEF above and beyond IQ and vice versa (per SNP). We chose mt-COJO to maximize sample size of the cEF summary statistics (i.e. the analysis could be run in the full sample). We then ran the same FUMA/MAGMA pipeline on the resulting summary statistics to discover what biological pathways remain after accounting for the other cognitive ability.

**Heritability and genetic correlations.**

To calculate cEF univariate heritability, we used BOLT-REML with a single variance component and considering all variants simultaneously^19^. We also used bivariate BOLT-REML to calculate the genetic correlation between cEF and IQ in the UKB.

We calculated genetic correlations of cEF with psychiatric, personality, neurological, and health related outcomes via LD Hub^20^ with the GWAS summary statistics from the full sample. LD Hub^20^ is a database of publicly available GWAS summary statistics and automated pipeline of LD score regression analysis that is utilized to estimate heritability of a trait of interest and genetic correlations with that trait and other relevant traits. We tested whether genetic correlations were significantly lower than 1.0 using a standard error calculated from the pseudovalues from the block jackknife output by LD score regression^10^.

We used GenomicSEM^21^ to estimate genetic relations of the UKB cognitive factor scores to a *p* factor predicting psychiatric diagnoses with available GWAS data. We included disorders that have been previously included in *p* factor models^22,23^: schizophrenia, bipolar disorder, major depressive disorder, anxiety, ADHD, alcohol use disorder, and eating disorder. Though summary statistics were available for obsessive-compulsive disorder and post-traumatic stress disorder, we did not include them because estimates would be underpowered given their small sample sizes (*n*s= 9,725 and 9,954 European ancestry). We did not include autism spectrum disorder or Tourette Syndrome, because they are not typically included in *p*-factor models. To improve fit, we included a residual correlation between bipolar disorder and schizophrenia; given the particularly high genetic correlation between these two disorders, it is reasonable to posit that they may jointly tap some unique variance (e.g. genetic variance specific to thought disorders). We used this *p*-factor model in correlational and multiple regression models with the UKB factor scores.

GenomicSEM conducts analyses in two stages. First, GWAS summary statistics are used to estimate the genetic covariance matrix and it sampling covariance matrix. Then these matrices are used to estimate the SEM using the lavaan package in R. GenomicSEM provides the following fit statistics: chi-square, CFI, and the standardized root mean square residual (SRMR). A nonsignificant chi-square, CFI > .95, and SRMR < .08 indicate good fit. CFI > .90 is sometimes considered to indicate acceptable fit^21^.

We also used GenomicSEM to estimate a series of multiple regressions of each trait on cEF, IQ, and Speed factor scores. The resulting betas indicate which cognitive factors genetically predict each trait, controlling for the genetic correlations among these factor scores. This analysis showed a pattern similar to the latent factor model.

**Polygenic score (PGS) analyses.**

To further validate the cEF phenotype in UKB, we used the UKB GWAS summary statistics to calculate PGSs for cEF and IQ (IQ) in two twin samples (the Colorado Longitudinal Twin study [LTS] and the Colorado Community Twin Sample [CTS])^24^. We then tested whether these PGSs differentially predicted EF latent variables and full-scale IQ in line with our conceptual model.

**Creation of PGSs in the CTS and LTS.** LTS and CTS samples were genotyped on the Axiom Precision Medicine Research Array 2.0 (Thermo-Fisher Scientific, Waltham, MA, USA). Genotypes were called using the APT software and Applied Biosystems Axiom Genotyping Solution Data Analysis Guide (publication Number 702961, https://assets.thermofisher.com/TFS-Assets/LSG/manuals/axiom_genotyping_solution_analysis_guide.pdf) and imputed to 1000 Genomes European reference panel via the Michigan Imputation Server^3^. When only one MZ twin was genotyped, scores for the other were duplicated for the PGS analysis. Post-imputation quality control (QC) included removing indels, rare-variants (MAF < 0.01), and variants that were poorly imputed (INFO < 0.3). Additional QC included removing variants with a high missing call rate, removing individuals with high missingness across the genome, removing rare variants, and removing variants out of Hardy-Weinberg equilibrium (--geno 0.02; --mind 0.02; --maf 0.01; --hwe 0.0001) before merging together datasets. After merging datasets together, another round of QC was done to make sure variants matched across datasets (--geno 0.02, --mind 0.02, --maf 0.01, --hwe 0.0001). PC analysis was completed to determine which individuals were of EUR ancestry, and then a second round of PC analysis was completed within the EUR ancestry individuals for inclusion as covariates in further analyses.

Next, summary best linear unbiased predictor (SBLUP) analyses were run to generate PGS for both cEF and IQ. Analyses employed all SNP summary statistics from the aforementioned UKB GWASs (i.e., not just those that met genome-wide significance [i.e., *p* < 5x10^-8^]), as restricting to genome-wide significant SNPs would bias the findings and poorly reflect the polygenic nature of these phenotypes. Effect sizes from UKB summary statistics were used to develop LD-adjusted effect sizes for all variants in common between our LD reference file (based on individuals of European Ancestry from the 1000 Genomes) and our CTS and LTS samples. SBLUP-derived PGSs were calculated using the --score function in PLINK (version 1.90b4.4).

**EF and IQ data in LTS and CTS.** We had two waves of EF data in the LTS, and one wave of EF data in the CTS. LTS twins completed a battery of 9 laboratory EF tasks at mean age 17 (*n* with genetic data = 755) and a modified battery of the same tasks again at age 23 (*n* with genetic data = 720). Methods and analyses for these tasks were fully described by Friedman et al. (2016)^7^. CTS twins (*n* = 563 with genetic data) completed identical versions as LTS wave 1 tasks at an average age 21 in a study that was run contemporaneously with the LTS wave 1 study; methods and results for the CTS sample were fully described by Friedman et al. (2020)^5^. Full scale IQ was measured in the LTS at age 16 with the Wechsler Adult IQ Scale (WAIS III [11 subtests]^25^; *n* = 776 with genetic data), and in the CTS at the same session as the EF testing with the Wechsler Abbreviated Scale of IQ (WASI [4 subtests]^26^; *n* = 561 with genetic data).

**PGS analysis model.** The 9 EF tasks at each wave included 3 response inhibition (antisaccade, stop-signal, and color-word Stroop), 3 working memory updating (keep track, letter memory, and spatial *n*-back), and 3 mental set-shifting (number–letter, color–shape, and category-switch) tasks, which were used to estimate the Unity/Diversity EF model^27^. In this model, a Common EF factor predicts all 9 tasks, and orthogonal Updating-specific and Shifting-specific factors also predict the updating and shifting tasks, respectively. As Common EF explains all the correlations among the response inhibition tasks, there is no inhibiting-specific factor.

To maximize power and minimize the number of tests, we created the model shown in Figure 3. This model was informed by our published twin models of these data^7^, which indicated high heritability of all three EF latent variables at both waves and in both samples (79% to 100%), with no new genetic influences in LTS across waves (*r*Gs = .99 to 1.0). Age, sex, and age*sex were regressed out of each task within each sample at each wave to obtain unstandardized residuals (using all available phenotypic data, not just those from people with genetic data). As the 9 EF tasks at wave 1 for the LTS and CTS were identical, they analyzed together, following our prior work^28^. The 9 EF tasks at wave 2 were included as additional EF factors (on which CTS participants were missing). Our prior work^7^ in the LTS found that the wave 2 Common EF, Updating-specific, and Shifting-specific factors showed identical genetic variance as the wave 1 factors, but new environmental influences on Common EF. Therefore, we modeled the 2 waves of EF data with Cholesky decompositions and regressed the first Cholesky factor for each EF latent variable on both the cEF and IQ PGSs, as well as the first 10 PCs and genotyping batch (3 dummy codes). As both the WAIS and WASI yielded scaled full-scale IQ scores, we age-and sex-regressed them within sample and analyzed them together in a single IQ variable, which was regressed on both the cEF and IQ PGSs, as well as the first 10 PCs and genotyping batch. Residual correlations were allowed between IQ and the first Cholesky factor for each EF, as well as between residuals for the same task across time.

Analyses were conducted in Mplus 8.3^29^, using the TYPE=COMPLEX option to cluster data by family. This option uses a weighted likelihood function and a sandwich estimator to obtain a scaled chi-square (χ^2^) and standard errors corrected for non-independence; prior studies demonstrate that it adequately corrects for nonindependence of twin data^30^. In the primary analysis, we restricted PGS analysis to individuals of European ancestry based on their scores on the first 3 PCs resulting in a final *n* of 916. Results for all individuals with genetic data, as well as individuals who with European ancestry based on the first 2 PCs are shown in the supplemental materials (supplemental Table S11). Because these analyses were intended to confirm that these factors in the UKB correspond to the factors in prior twin studies that have motivated this GWAS approach, these are truly *a priori* hypotheses and we use a standard alpha of .05, uncorrected.

**Annotation through FUMA/MAGMA pipeline.**

Using data from the 1000 Genomes Project (1000G)^31^ phase 3 European (EUR) population as a reference, LD structure (r^2^) of pairwise SNPs and minor allele frequencies (MAFs) were pre-computed. Independent significant SNPs (r^2^ < 0.6) having a genome-wide significant *p*-value (*p* < 5 x 10^-8^) were distinguished. All SNPs available in the 1000G^31^ EUR reference panel, in LD with the independent significant SNPs (r^2^ ≥ 0.6, max 1Mb window, MAF ≥ 0.01) were defined as candidate SNPs for association with cEF. Of the identified significant SNPs, those independent at r^2^ < 0.1 were defined as lead SNPs. LD blocks of all the identified independent significant SNPs and lead SNPs that were less than 250 kb apart were combined and characterized as genomic risk loci. Because the power of GWAS depends upon how well causal variants are tagged, annotation is extended beyond independent significant SNPs to incorporate all candidate SNPs. Thus, candidate SNPs are functionally annotated and used for our gene prioritization analyses, while lead SNPs with the lowest *p*-value are used to represent their respective genomic loci.

**Functional characterization of lead independent SNPs.** To determine the functional consequences of SNPs significantly associated with cEF, ANNOVAR^32^ was run on candidate SNPs located within the independent genomic loci to determine their functional consequences in FUMA (defaults: r^2^ ≥ 0.6, *p* < 0.05, MAF ≥ 0.01). These SNPs were matched according to chromosomal location, base pair position, reference and non-reference alleles and then annotated accordingly. To map candidate SNPs significantly associated with cEF to genes, two different strategies were applied based on Ensembl genes (build 85) using FUMA^15^. First, SNPs on or near genes, determined via ANNOVAR^32^ annotation, were positionally mapped to genes based on their physical distance (< 10 kb) from protein-coding genes.

Second, to determine if significantly associated SNPs related to gene expression, SNPs were further annotated by FUMA expression quantitative trait loci (eQTLs) status. SNPs that significantly affect gene expression were extracted from the GTEx sample^33^ and BRAINEAC^34^ sample. SNPs were mapped to genes within a 1 Mb window, known as cis-eQTLs, and were limited to only significant SNP-gene pairs (false discovery rate [FDR] ≤ 0.05, the default in FUMA). GTEx v7 Brain and BRAINEAC database of brain-tissue-specific gene expression data were utilized to perform eQTL mapping of the following brain tissues: amygdala, anterior cingulate cortex BA24, caudate basal ganglia, cerebellar hemisphere, cerebellum, cortex, frontal cortex BA9, hippocampus, hypothalamus, nucleus accumbens basal ganglia, putamen basal ganglia, cervical (c-1) spinal cord, and substantia nigra. BRAINEAC eQTLs were used for the following brain tissues: cerebellar cortex, frontal cortex, hippocampus, inferior olivary nucleus, occipital cortex, putamen, substantia nigra, temporal cortex, thalamus, and intralobular white matter.

**Regulatory elements of intronic SNPs: CADD and regulome scoring and 3D chromatin interaction.** To determine whether intronic independent SNPs served a possible regulatory functioning, we annotated significant independent SNPs for their Combined Annotation Dependent Depletion score (CADD), Regulome score and for chromatin-chromatin interaction via 3D chromatin interaction (Hi-C)^35^. CADD scoring shows the likelihood that the SNP is a deleterious mutation. Regulome scores SNPs based on their likelihood the intronic SNPs have cis-regulatory function. Hi-C examines whether SNPs represent long-range enhancer-promotor associations. For Hi-C data from the following pre-existing builds were used in FUMA: dorsolateral PFC, hippocampus, and neural progenitor cells.

**Gene-level regression analyses via MAGMA.**

**Gene-based analysis.** To determine what genes are significantly associated with cEF and create a prioritized list of genes based on the degree of association, MAGMA^36^ gene analysis was performed in FUMA. MAGMA Uses a multiple regression model run on GWAS summary statistics designed to incorporate LD between genetic variants and detect the aggregated effects of multiple weakly associated variants. MAGMA combines the *p*-values of SNPs, mapped to protein-coding genes, to generate a gene-based *p*-value, in addition to genetic correlations between neighboring genes. This analysis produces a prioritized list of significantly associated, protein-coding genes to quantify the level of association between the identified genes and EF. We ran this analysis in the densely and sparsely phenotyped samples, as well as the full sample.

**Gene-set analysis.** To detect biological pathways significantly associated with cEF, we used MAGMA to run a competitive gene-set analysis and cell-type specific gene-set analysis. This analysis also accounts for potential confounding variables, such as gene density and size^37^. A competitive gene-set analysis is a gene-level linear regression model designed to determine whether genes within a gene-set have a significantly greater association with cEF than all other genes outside of the gene set. Gene sets are determined by shared biological and functional characteristics between genes defined by the datasets in MBsig6.1^38^. For the cell-type specific analysis, we annotated our findings with QTL information from RNA cell-type specific studies of human postmortem cortex^39^, hippocampus^40^, and frontal cortex^41^ (during prenatal development).

**Gene-property analysis to determine tissue specificity.** To evaluate in what tissues SNP effects across the whole genome are likely expressed, MAGMA gene-property analysis was performed in FUMA. This gene-property analysis was performed on 30 general and 53 specific GTEx v7^33^ tissue types.

**Transcription patterns.**

**TWAS with PrediXcan.** To identify genetic transcription patterns implicated by the whole-genome SNP effects and eQTL results, we ran a Transcriptome-Wide Analysis (TWAS) using PrediXcan^17^. PrediXcan imputes gene expression from SNPs via an elastic net model trained in an eQTL sample, in this case the GTeX sample^33^. We chose PrediXcan because it uses summary statistics and allows variability across tissue types^42^. We ran PrediXcan with the summary statistics for our cEF GWAS separately for each tissue that was significant by MAGMA gene-property analysis.

**Predicted Transcription-Based Drug Repurposing.** Because TWAS offers a predicted transcriptional profile, this profile can be compared to other datasets from computational pharmacogenomics. So et al. (2017)^43^ expanded on PrediXcan TWAS using the connectivity Map (cMAP) Library^44^ to infer what drugs mimic the implicated transcription pattern. The cMAP is a dataset of experimental transcription changes in stem cell lines after exposure to a pharmaceutical substance^44^. We entered all significant inferred transcripts post-Bonferroni correction and their predicted differential expression *z*-scores into the Connectivity map toolbox to obtain the top 15 substances predicted to reverse the pattern of transcription^43^.
